# Mechanism of RPA-Facilitated Processive DNA Unwinding by the Eukaryotic CMG Helicase

**DOI:** 10.1101/796003

**Authors:** Hazal B. Kose, Sherry Xie, George Cameron, Melania S. Strycharska, Hasan Yardimci

## Abstract

The DNA double helix is unwound by the Cdc45/Mcm2-7/GINS (CMG) complex at the eukaryotic replication fork. While isolated CMG unwinds duplex DNA very slowly, its fork unwinding rate is stimulated by an order of magnitude by single-stranded DNA binding protein, RPA. However, the molecular mechanism by which RPA enhances CMG helicase activity remained elusive. Here, we demonstrate that engagement of CMG with parental double-stranded DNA (dsDNA) at the replication fork impairs its helicase activity, explaining the slow DNA unwinding by isolated CMG. Using single-molecule and ensemble biochemistry, we show that binding of RPA to the excluded DNA strand prevents duplex engagement by the helicase and speeds up CMG-mediated DNA unwinding. When stalled due to dsDNA interaction, DNA rezipping-induced helicase backtracking re-establishes productive helicase-fork engagement underscoring the significance of plasticity in helicase action. Together, our results elucidate the dynamics of CMG at the replication fork and reveal how other replisome components can mediate proper DNA engagement by the replicative helicase to achieve efficient fork progression.

## Introduction

All cells utilize a ring-shaped helicase that separates the two strands of the DNA double helix during chromosome replication. Replicative helicases form a homohexameric complex such as gp4 in bacteriophage T7, DnaB in bacteria, large T antigen in Simian Virus 40 (SV40), E1 helicase in bovine papillomavirus, and minichromosome maintenance (MCM) in archaea (Trakselis, 2016). The only known exception to the homohexameric nature is the eukaryotic replicative DNA helicase, comprising the Mcm2-7 motor containing six different but highly related AAA+ ATPases. In G1 phase of the cell cycle, Mcm2-7 rings are loaded onto duplex DNA as double hexamers (Abid Ali et al., 2017; Evrin et al., 2009; Noguchi et al., 2017; Remus et al., 2009). Activation of the helicase in S phase occurs upon binding of Cdc45 and GINS, and subsequent remodeling of Mcm2-7 from encircling double-stranded (ds) to single-stranded (ss) DNA in its central channel (Douglas et al., 2018; Fu et al., 2011; Kose et al., 2019). Through ATP hydrolysis, Cdc45/Mcm2-7/GINS (CMG) complex translocates along the leading-strand template in the 3ʹ to 5ʹ direction and unwinds DNA at the replication fork (Fu et al., 2011). In addition to its role in DNA unwinding, the replicative helicase acts as a hub to organize other replication factors around itself, thus assembling the replisome.

CMG was first characterized biochemically in isolation by purifying the complex from *Drosophila* embryo extracts (Moyer et al., 2006). Recombinant *Drosophila*, yeast and human CMG complexes were later shown to unwind DNA substrates containing Y-shaped fork structures (fork DNA) in an ATP-dependent manner (Ilves et al., 2010; Kang et al., 2012; Langston et al., 2014). When unwinding DNA at the fork, isolated CMG can freely bypass protein obstacles on the lagging-strand template indicating that this strand is excluded from the helicase central channel during translocation (Kose et al., 2019). Thus, CMG functions via steric exclusion, a mechanism shared by all known replicative helicases (Egelman et al., 1995; Fu et al., 2011; Kaplan, 2000; Lee et al., 2014; Yardimci et al., 2012b).

Using single-molecule magnetic-tweezers, we earlier found that individual *Drosophila* CMG complexes exhibit forward and backward motion while unwinding dsDNA (Burnham et al., 2019), similar to E1 and T7 gp4 helicases (Johnson et al., 2007; Lee et al., 2014; Syed et al., 2014). Furthermore, the helicase often enters long-lived paused states leading to an average unwinding rate of 0.1-0.5 base pairs per second (bps^−1^), which is approximately two orders of magnitude slower than eukaryotic replication fork rates observed *in vivo* (Anglana et al., 2003; Raghuraman et al., 2001). However, recent single-molecule work with yeast CMG suggests that the helicase translocates on ssDNA at 5-10 bps^-1^ (Wasserman et al., 2019). Single-molecule trajectories by other replicative helicases such as DnaB and gp4 suggest that helicase pausing during dsDNA unwinding is a general property of these enzymes (Ribeck et al., 2010; Syed et al., 2014). However, it is not clear why replicative helicases frequently halt whilst moving at the fork and how higher speeds are achieved by the entire replisome.

The rate of DNA unwinding by *E.coli* DnaB and T7 gp4 is substantially enhanced when engaged with their corresponding replicative polymerases (Kim et al., 1996; Stano et al., 2005) suggesting that rate of fork progression in eukaryotes may also depend on DNA synthesis. Likewise, uncoupling of CMG from the leading-strand polymerase leads to fork slowing in an *in vitro* purified yeast system (Taylor and Yeeles, 2019). 5-10 fold reduction in helicase speed was also observed in *Xenopus* egg extracts when DNA synthesis was inhibited by aphidicolin (Sparks et al., 2019). However, this decrease in CMG translocation rate is not sufficient to account for the approximate 100-fold lower rates seen in DNA unwinding by isolated CMG (Burnham et al., 2019). Thus, in addition to polymerases, other replisome associated factors may be essential to increase the rate of unwinding by the helicase. Intriguingly, single-molecule visualization of the ssDNA-binding protein RPA during CMG-driven DNA unwinding indicated that *Drosophila* CMG proceeds at an average rate of 8 bps^−1^ at the fork. This result suggests that binding of RPA to unwound DNA improves the rate of translocation by CMG. One possible explanation for RPA-induced rate increase is the association of RPA with the translocation strand (the leading-strand template) behind CMG and concomitant hindrance of helicase backtracking. In addition, coating of the lagging-strand template by RPA may prevent rezipping of the two strands in front of the helicase, thus increasing the rate of unwinding. Finally, RPA binding may also influence helicase activity by altering the interaction of CMG with the excluded strand.

Control of DNA unwinding by replicative helicases through their interaction with the excluded strand has been demonstrated in different organisms. For example, while wrapping of the displaced strand around an archaeal MCM was proposed to increase its helicase activity (Graham et al., 2011), interaction of DnaB with the displaced strand through its exterior surface adversely affects DNA unwinding (Carney et al., 2017). Although, it is not clear whether CMG makes contacts with the lagging-strand template via specific residues on its outer surface, the presence of the excluded strand is important for unwinding of dsDNA by CMG. Notably, unwinding of synthetic DNA substrates by CMG relies not only on the availability of a 3ʹ ssDNA tail for CMG binding, but also on the presence of a 5ʹ overhang. On partially duplexed DNA lacking the 5ʹ flap, CMG binds the 3ʹ ssDNA and subsequently slides onto dsDNA upon meeting the duplexed region (Kang et al., 2012; Langston and O’Donnell, 2017). Equally, T7 gp4, DnaB, and archaeal MCM can transfer from translocating on ssDNA to dsDNA without unwinding the template when encountering a flush ss-dsDNA junction (Jeong et al., 2013; Kaplan, 2000; Kaplan et al., 2003; Shin et al., 2003). This unproductive translocation on duplex DNA is likely a consequence of the central pores of these motors being sufficiently large to accommodate dsDNA.

In this study, we sought to address the mechanism by which RPA stimulates DNA unwinding rate of the eukaryotic CMG helicase. Using ensemble and single-molecule assays, we show that the helicase can stall when interacting with the replication fork due to partial entry of the parental duplex into the Mcm2-7 pore. Our results suggest that DNA rewinding-induced reverse helicase movement can rescue the CMG from this paused conformation highlighting the advantage of plasticity in helicase motion. Importantly, RPA binding to the excluded strand impedes helicase engagement with duplex DNA and stimulates unwinding, which reveals an important role for RPA in regulating replisome progression.

## Results

### Direct visualization of RPA-facilitated processive fork unwinding by individual CMG molecules

We previously demonstrated CMG-driven unwinding of surface-immobilized 2.7-kb dsDNA substrates through accumulation of EFGP-tagged RPA (EGFP-RPA) using total internal reflection fluorescence (TIRF) microscopy (Kose et al., 2019). To more directly assess the processivity and rate of DNA unwinding by single CMG molecules, we examined the translocation of fluorescently-labelled CMG complex on DNA molecules stretched along the glass surface of a microfluidic flow cell. A 10-kb fragment of λ DNA was ligated to a short fork DNA substrate at one end and to a digoxigenin-modified DNA fragment on the opposite end. The 5ʹ tail of the forked end contained a biotin to attach the 10-kb substrate to biotin-functionalized glass through biotin-streptavidin binding. The digoxigenin-modified end was coupled to anti-digoxigenin-coated microsphere to stretch DNA molecules by buffer flow, and to subsequently attach this end to the surface (Figure 1a, details are described in the Methods section). The 3ʹ tail of the fork contained a 40-nt polyT ssDNA (dT_40_) for CMG binding and a Cy3 fluorophore to follow the position of the translocation strand (Figure 1a). After immobilizing DNA on the surface (Figure 1b), CMG that was labelled with LD655 dye on Mcm3 (CMG^LD655^) was drawn into the flow cell and incubated in the presence of ATPγS. Subsequently, the CMG-ATPγS mixture in the flow channel was exchanged with a solution containing ATP and EGFP-RPA (Figure 1c). Near-TIRF imaging was performed in the absence of buffer flow through the flow cell. We observed EGFP-RPA binding initially at the forked end of the stretched DNA before growing as a linear tract towards the microsphere-tethered end (Figure 1d, left panel). The leading-strand template bound by EGFP-RPA appeared as a compact diffraction-limited spot moving at the fork because this strand was not attached to the surface. When either CMG from ATPγS-containing buffer or ATP from subsequent RPA-supplemented solution were omitted, we did not detect linear EGFP-RPA tracts (data not shown) indicating that we are visualizing CMG-dependent fork unwinding. Approximately 25% of the growing RPA tracts contained labelled CMG translocating at the fork (Figure 1d, middle panel). The relatively low fraction of labelled CMG molecules is most likely due to inefficient conjugation of the fluorophore or subsequent photobleaching.

**Figure 1.**
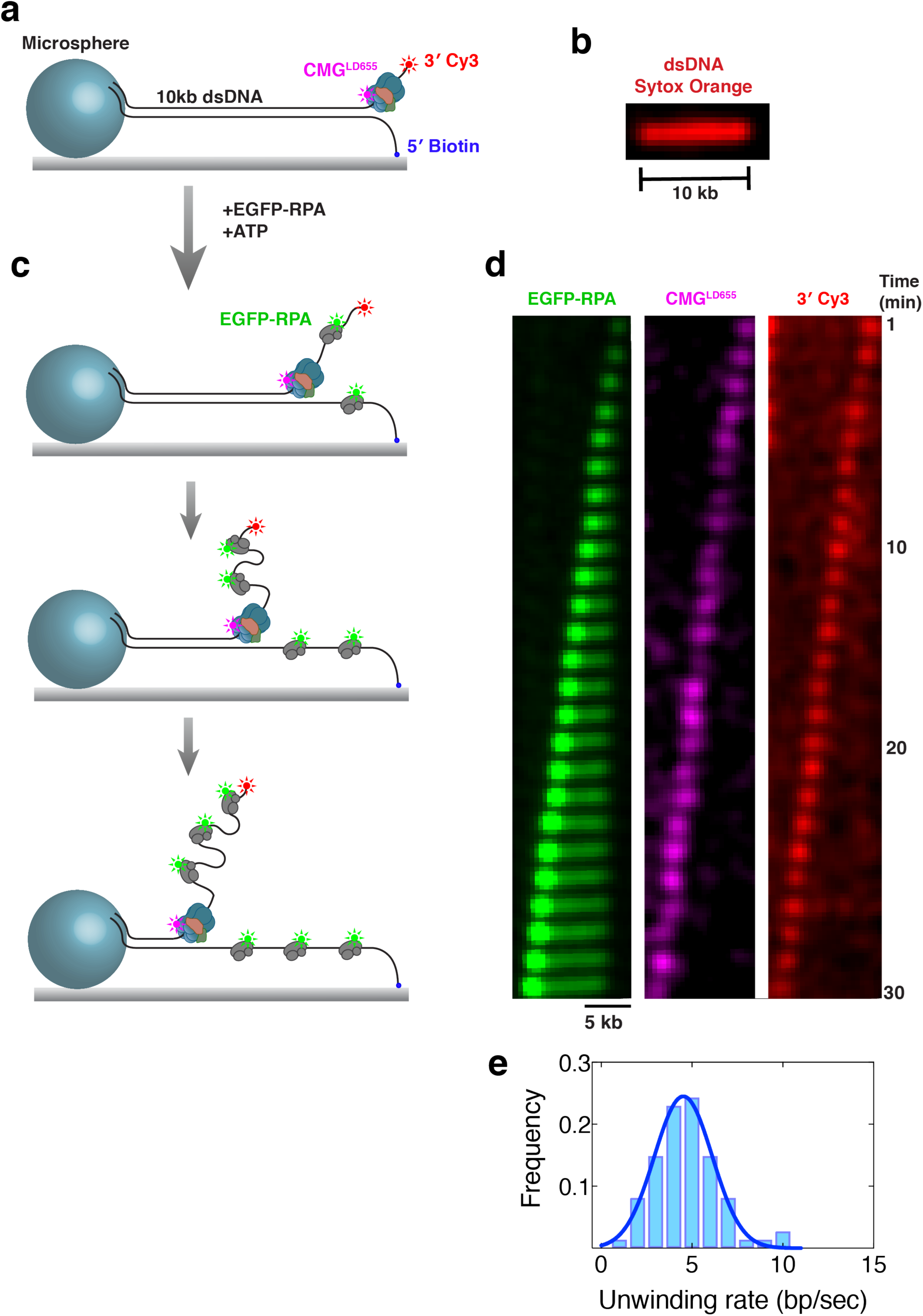
Direct visualization of RPA-facilitated processive fork unwinding by individual CMG molecules. **a,** A cartoon illustration of 10-kb linear DNA with forked end attached to the surface, and the opposite end bound to a surface-immobilized microsphere. The DNA substrate was labelled with Cy3 at the 3’ dT_40_ ssDNA tail near the fork junction. LD655-labelled CMG was bound to the dT_40_ ssDNA in the presence of ATPγS. **b,** A sample stretched 10-kb linear DNA stained with a fluorescent dsDNA intercalator Sytox Orange. **c,** After binding CMG on the surface-immobilized DNA, EGFP-RPA and ATP was introduced to initiate unwinding. While CMG unwinds DNA at the fork, EGFP-RPA binds both strands of unwound DNA. **d,** Kymograph showing a representative 10-kb DNA being unwound entirely by a single CMG complex. EGFP-RPA (left panel), LD655-labelled CMG (center panel), and 3’ Cy3 (right panel) are imaged during unwinding under near-TIRF conditions. Images were acquired in the absence of buffer flow. **e,** Histogram of CMG-catalyzed DNA unwinding rates on stretched DNA.

The Cy3-labelled translocation strand co-localized with the bright EGFP-RPA spot leading in front of EGFP-RPA tract as expected (Figure 1d, right panel). Many 10-kb DNA molecules were unwound entirely at an average rate of 4.5 ± 1.6 bps^-1^ (Figure 1e). The majority of molecules were not fully unwound, likely being interrupted because of CMG encountering nicks present on stretched DNA. On some molecules, unwound region of the RPA-coated leading-strand template either uncoupled from (Supplementary Figure 1a) or diffused along (Supplementary Figure 1b) the stretched lagging-strand template. We attribute these observations to CMG encountering a nick on the leading-strand template. Labelled CMG always dissociated from DNA upon hitting a leading-strand nick suggesting that the helicase ran off the free 5ʹ end of the translocation strand (Supplementary Figures 1a and 1b). In other cases, partially unwound stretched DNA broke suggesting that CMG ran into a nick on the lagging-strand template (Supplementary Figure 1c). While the unwound lagging-strand template instantly moved to the point where it was tethered through the 5ʹ biotin, the unwound leading-strand template and CMG moved towards the microsphere (Supplementary Figure 1c). In our previous study (Kose et al., 2019), we used 2.7-kb DNA substrates tethered to the surface at one end and measured unwinding rates relying solely on the level of EGFP-signal intensity. Therefore, any event where CMG encountered a nick was scored as the entire DNA being unwound, which may have led to an overestimation of unwinding rate (8.2 bps^-1^ as opposed to 4.5 bps^-1^). Even though we were unable to measure the upper limit of CMG processivity, our results clearly demonstrate that when RPA is available, individual CMG helicases can unwind thousands of base pairs of dsDNA at a rate matching the ssDNA translocation rate of the helicase (Wasserman et al., 2019).

### RPA increases the rate of DNA unwinding by CMG

The rate of duplex unwinding by CMG, measured by magnetic tweezers assay (Burnham et al., 2019), was found to be strikingly low (0.1-0.5 bps^-1^) compared to that measured by single-molecule fluorescence imaging in the presence of fluorescent RPA (Figure 1). Therefore, the translocation speed of CMG that unwinds dsDNA at the fork must be stimulated by an order of magnitude by RPA. Because these measurements were done using two different experimental methods under different conditions, we wanted to compare the rate of CMG translocation at the fork with and without RPA using a single assay. To this end, we examined CMG helicase activity at the ensemble level on a fork DNA substrate containing 10 repeats of GGCA sequence, d(GGCA)_10_, on the 5ʹ lagging-strand arm, dT_40_ as the 3ʹ arm, and 236 bp of dsDNA (Figure 2a). CMG was first incubated with the fork substrate in the presence of ATPγS for its binding to the 3ʹ tail. The d(GGCA)_10_ sequence folds into secondary hairpin-like structures and prevents CMG binding to this strand (Petojevic et al., 2015). After CMG binding, ATP was added to the reaction to trigger helicase translocation. When RPA was added simultaneously with ATP, we observed extensive unwinding of the substrate in a CMG-dependent manner (Figure 2a, lane 4). In contrast, no unwinding was detected when RPA was omitted from the reaction (Figure 2a, lane 3), most likely due to re-annealing of the complementary strands behind the helicase. To overcome this limitation, we generated a similar fork substrate that contained a Cy5-labelled oligonucleotide downstream of a 252 bp duplex. The Cy5-labelled strand contained a d(GGCA)_10_ 5ʹ tail and 28-nt complementary sequence to the translocation strand (Figure 2b). Therefore, CMG is expected to displace the Cy5-modified strand if it can translocate through the 252-bp dsDNA even if DNA rewinds in the wake of the advancing helicase. To measure the kinetics of CMG translocation, the reaction was quenched at different times after ATP addition. When RPA was added together with ATP, CMG rapidly displaced the Cy5-labelled strand (Figure 2c). To quantify CMG-dependent unwinding, the data was corrected for Cy5-modified strand displaced by RPA alone (Supplementary Figure 2a). CMG displaced the Cy5-modified strand to a significant extent even in the absence of RPA (Figure 2d) indicating that lack of strand separation on the 236-bp duplex fork (Figure 2a) was due to rewinding of DNA trailing the helicase. Unwinding of the 28-bp duplex region was dependent on CMG binding to the upstream 3ʹ dT_40_ ssDNA tail (Supplementary Figure 2b) indicating that the helicase translocated through the 252-bp duplex before displacing the Cy5-modified strand. In the presence of RPA, CMG unwound all 280 bp (252 bp + 28 bp) at an observed rate of k_obs_ = 1.31 ± 0.11 min^-1^ (Figure 2e, solid line) leading to an average DNA unwinding rate of 6.1 ± 0.5 bps^-1^ (280 bp × k_obs_), in good agreement with single-molecule measurements (Figure 1). In contrast, CMG alone displaced the Cy5-modified strand at a rate of k_obs_ = 0.048 ± 0.003 min^-1^ indicating an average DNA unwinding rate of 0.22 ± 0.01 bps^-1^ (Figure 2f, solid line), consistent with linear unwinding rates measured with magnetic tweezers (Burnham et al., 2019). Therefore, the gel-based helicase assays described here suggest that RPA increases the rate of CMG translocation at the fork about an order of magnitude, in agreement with single-molecule studies.

**Figure 2.**
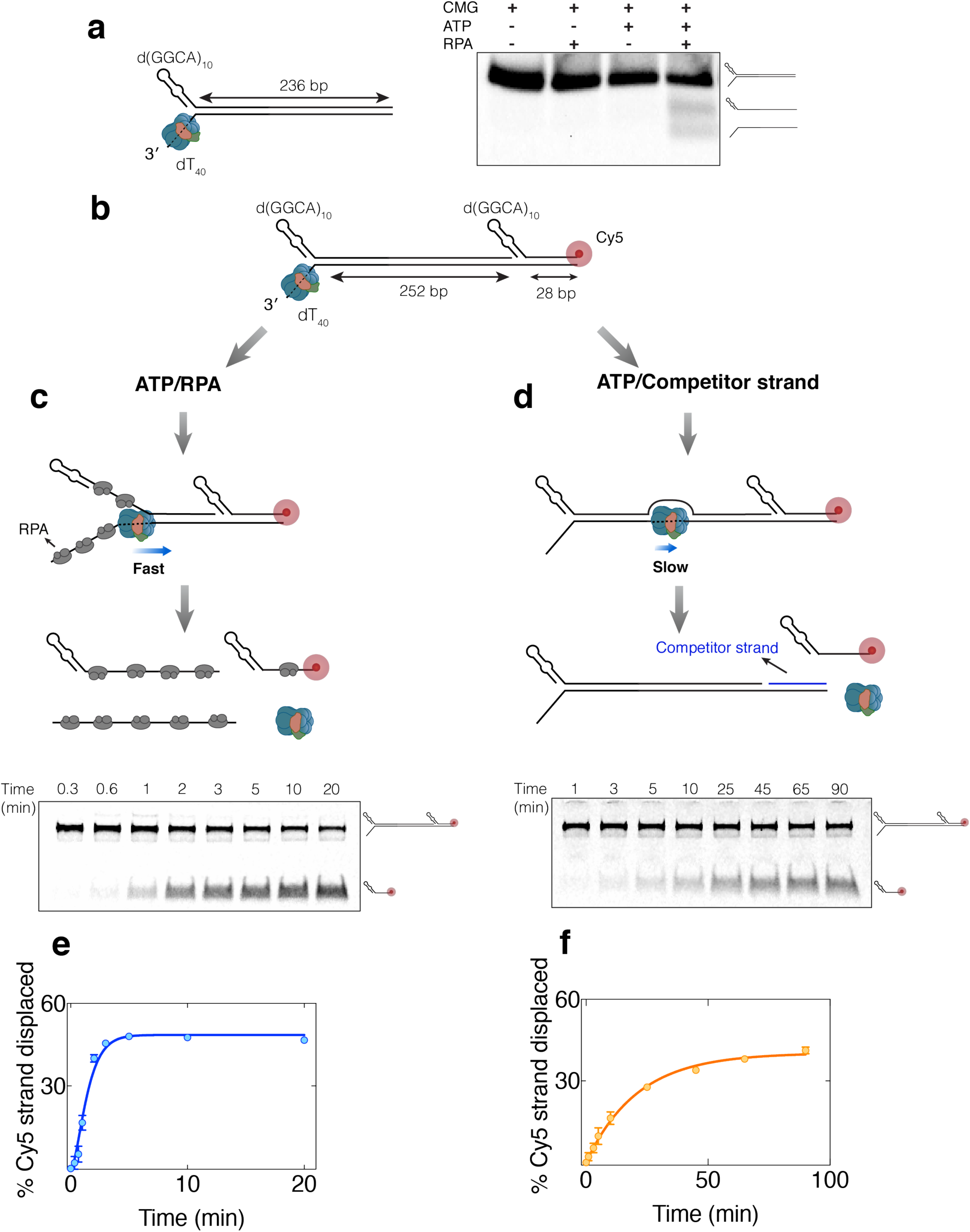
RPA increases the rate of DNA unwinding by CMG. **a,**Fork DNA containing 236-bp dsDNA was incubated with CMG in the presence of ATPγS for binding to 3ʹ dT_40_ tail. ATP was added with (lane 4) or without (lane 3) RPA and incubated further before separating on 3% agarose. DNA was labelled internally with multiple Cy5 fluorophores on both strands. **b,** Fork DNA substrate containing 252-bp long dsDNA, followed by 28-bp duplex and Cy5-modification on the excluded strand was bound by CMG. ATP was added to initiate translocation by the helicase. **c,** CMG-mediated displacement of Cy5-labelled strand in the presence of RPA. When included in ATP buffer, RPA prevents reannealing of DNA behind the helicase as well as new CMG binding. **d,** CMG-mediated displacement of Cy5-labelled strand in the absence of RPA. DNA rewinds within the 252-bp duplex region. To prevent rehybridization of the Cy5-labelled strand to long DNA substrate, excess competitor oligonucleotide containing complementary 28-nt sequence to long DNA was added with ATP. To achieve single-turnover kinetics, excess dT_40_ oligonucleotide was included in ATP buffer that captures any free CMG. **e-f,** Percentage of Cy5-modified strand unwound versus time by CMG in the presence (**e**) and absence of RPA (**f**). The data represent mean ± SD (n=2), and were fit to Equation 1 with m = 1 or 2 (see Methods) resulting in best R-squared value. Solid lines are fits to Equations 3 and 2 in **e** and **f**, respectively.

### Duplex DNA engagement by CMG at the fork impedes its helicase activity

Our single-molecule magnetic tweezers measurements indicated that the slow nature of DNA unwinding by isolated CMG was due to frequent entry of the helicase into long-lived paused states at the fork (Burnham et al., 2019). We wished to determine the origin of these pausing events. Given the ability of CMG to encircle dsDNA, we considered the possibility that while translocating along the leading-strand template, the helicase may engage with the parental duplex at the fork junction as an off-pathway interaction. In fact, inhibition of helicase activity was reported for T7 gp4 due to its interaction with the parental dsDNA on fork DNA substrates (Jeong et al., 2013). In addition, enhanced DnaB helicase activity upon duplexing the leading-strand arm (excluded strand) of fork DNA has been attributed to a decrease in probability of the helicase encircling parental duplex DNA at the fork (Atkinson et al., 2010; Kaplan, 2000). In support of this model, constricting the central channel of DnaB by point mutations led to elevated levels of fork DNA unwinding (Strycharska et al. 2013). Likewise, while unwinding dsDNA, CMG may frequently slip onto the fork nexus to capture a fragment of duplex DNA in its central channel and potentially enter into a non-translocating state. Based on this model, duplexing and thus stiffening and enlarging the displaced arm of fork DNA should stimulate DNA unwinding by impeding the helicase ring encircling duplex DNA at the fork junction, as seen for DnaB (Kaplan, 2000). In previous work, we used fork substrates with d(GGCA)_10_ on the 5ʹ lagging-strand arm (Kose et al., 2019). As this sequence folds into secondary structures, it may prevent CMG partially encircling the parental duplex at the fork junction. We found that replacing d(GGCA)_10_ on the 5ʹ tail with dT_40_ led to two-fold decrease in DNA unwinding by CMG (Supplementary Figure 3a) in line with the prediction that duplex engagement at the fork junction impairs CMG translocation. We tested this model in more detail using a fluorescence-based single turn-over kinetic unwinding assay (Kose et al., 2019). The fork DNA templates used in this assay contained 28-bp duplex DNA, a dT_40_ leading-strand arm and 22-nt long either single-stranded (Fork^ssLag^) or duplexed lagging-strand arm (Fork^dsLag^) (Figure 3a and Supplementary Figure 3b). Fork substrates were modified with Cy5 at the 5ʹ end of the leading-strand template and a BHQ2 quencher on the complementary strand so that strand separation leads to increase of Cy5 fluorescence (Figure 3a). To detect single turn-over unwinding kinetics, CMG was pre-bound to the fork in the presence of ATPγS, and subsequently ATP was added together with a competitor oligonucleotide to sequester free CMG (Supplementary Figure 3c). We confirmed that the fluorescence increase was dependent on CMG pre-binding as well as subsequent addition of ATP on the 28-bp duplex fork (Supplementary Figure 3d). Importantly, Fork^dsLag^ showed higher fluorescence increase than Fork^ssLag^ (Figure 3b) indicating a greater level of unwinding consistent with our hypothesis. To ensure this effect is not due to a discrepancy in the efficiency of helicase binding the two substrates, CMG was first bound to Fork^ssLag^ before triggering unwinding with ATP in the presence of an additional oligonucleotide complementary to the lagging-strand arm (Comp^Lag^). The addition of Comp^Lag^ with ATP was sufficient to increase the level of fork unwinding (Supplementary Figure 3e) demonstrating that duplexing the 5ʹ tail stimulates CMG helicase activity rather than fork binding.

**Figure 3.**
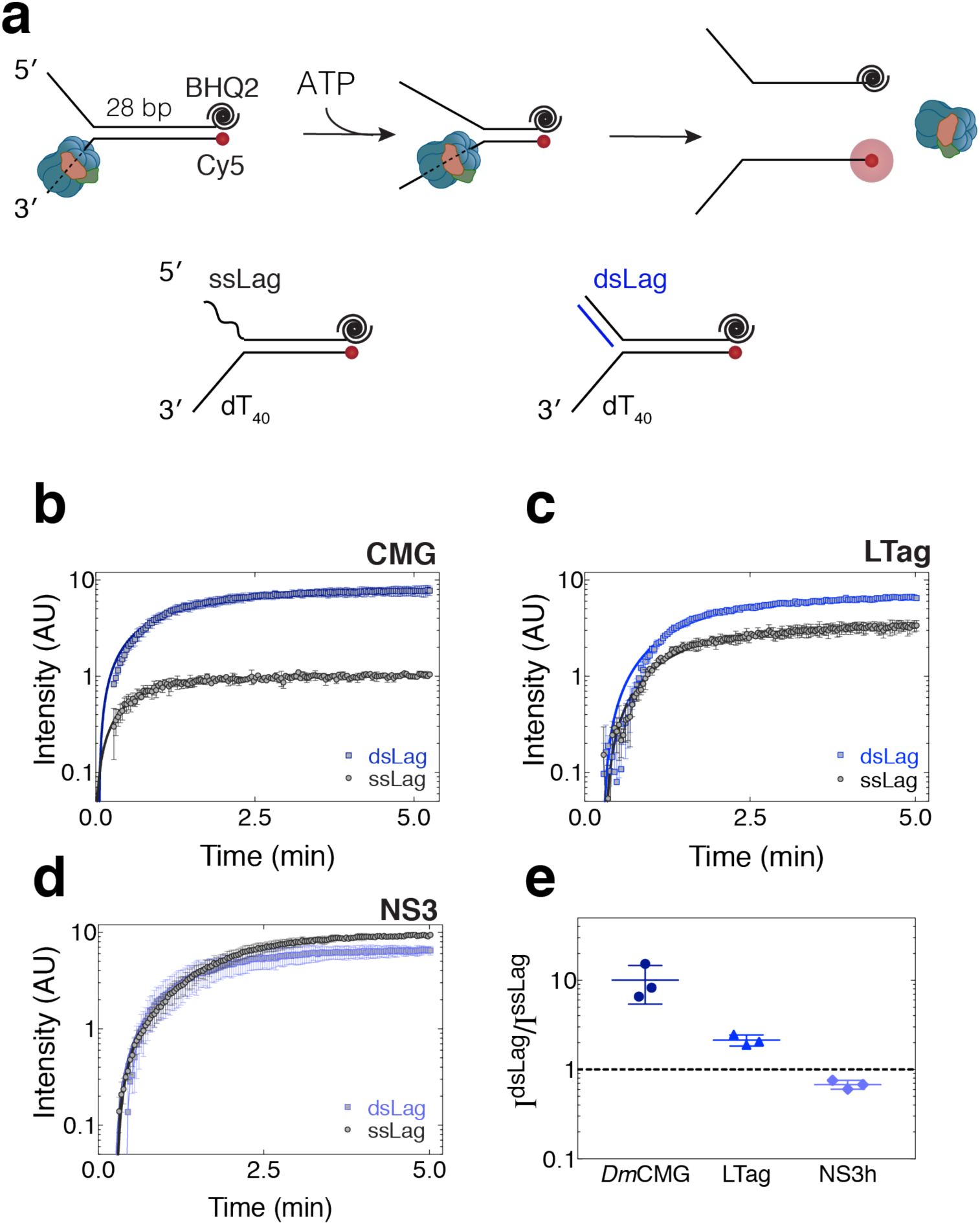
Duplex DNA engagement by CMG at the fork impedes its helicase activity. **a-d**, Single turn-over unwinding of fork DNA substrates containing ss (black) or ds (blue) lagging-strand arm by CMG (**b**), large T antigen (**c**), and NS3 (**d**). DNA substrates contained 28-bp duplex region and were modified with Black Hole Quencher 2 (BHQ2) at the 3ʹ end of the lagging-strand template and Cy5 at the 5ʹ end of the leading-strand template. Solid lines represent fits to Equation 2 in **b** and Equation 3 in **c** and **d** (see Methods). **e,** The ratio of fluorescence plateau intensity on Fork^dsLag^ to Fork^ssLag^ indicates that DNA stimulation of fork DNA unwinding by duplexing the lagging-strand arm is helicase dependent. All data represent mean ± SD (n=3).

To rule out the possibility that improved unwinding of Fork^dsLag^ is an inherent feature of the DNA template and not the CMG helicase, we analyzed unwinding kinetics of the same fork substrates by large T antigen and NS3, both 3ʹ-to-5ʹ helicases. Although large T antigen hexamers are proposed to encircle dsDNA at the origin of replication (Gai et al., 2016), the helicase central channel within the motor domain is narrower than the cross section of dsDNA. As a result, large T antigen can unwind partial duplex DNA substrates lacking a 5ʹ-ssDNA flap (Wiekowski et al., 1988). Because large T antigen should be less susceptible to sliding onto duplex DNA at the fork, stimulation of helicase activity by duplexing the 5ʹ tail of the fork is expected to be less pronounced compared to CMG. Consistently, while fluorescence increase on Fork^dsLag^ was 10-fold higher than that of Fork^ssLag^ when unwound by CMG, this change was only 1.5-fold in large T antigen-catalyzed unwinding of the two substrates (Figures 3c and 3e). This result suggests that regulation of DNA unwinding by replicative helicases through duplex DNA interaction at the fork depends on the size of their central channel. To further test our model, we measured unwinding of fork DNA by NS3 from hepatitis C virus, a monomeric helicase lacking a central pore as found in replicative helicases (Gu and Rice, 2010; Pang et al., 2002). As expected, no improvement of NS3 helicase activity was detected when the 5ʹ tail of the fork was duplexed (Figures 3d and 3e). Together, our data strongly suggest that duplexing the lagging-strand arm of fork DNA stimulates CMG helicase activity by preventing the helicase pore from encircling duplex DNA at the fork junction.

### Single-molecule analysis of CMG-catalyzed fork unwinding

To independently validate the results from fluorescence-based ensemble experiments demonstrated in Figure 3, we devised a single-molecule assay to directly monitor DNA unwinding by CMG. To this end, fork DNA bearing a biotin at the 5ʹ end of the translocation strand and an Atto647N fluorophore at the 5ʹ end of the displaced strand was immobilized on the streptavidin-functionalized glass surface of a microfluidic flow cell (Figure 4a). CMG was introduced in the presence of ATPγS and allowed to bind the 3ʹ tail of surface-tethered fork DNA. Subsequently, ATP was drawn into the flow cell to initiate unwinding while the Atto647N-labelled strand was imaged with TIRF microscopy. Unwinding of the fork substrate was assessed from the loss of fluorescence signal upon dissociation of the displaced strand, which was coupled to the surface only through its hybridization with the biotin-modified translocation strand. After correcting for photobleaching (Supplementary Figure 4), we found that 20-30% of surface-immobilized DNA was unwound by CMG (Figures 4a and 4c, Fork^ssLag^). To examine whether duplexing the lagging-strand arm of the fork alters the efficiency of unwinding, we annealed a complementary oligonucleotide to the 5ʹ ssDNA tail of immobilized fork DNA before CMG binding. Upon addition of ATP, 60-70% of Fork^dsLag^ was unwound (Figures 4b and 4c, Fork^dsLag^). Therefore, single-molecule visualization of DNA unwinding confirms that duplexing the lagging-strand tail of fork DNA has a profound stimulatory effect on CMG helicase activity.

**Figure 4.**
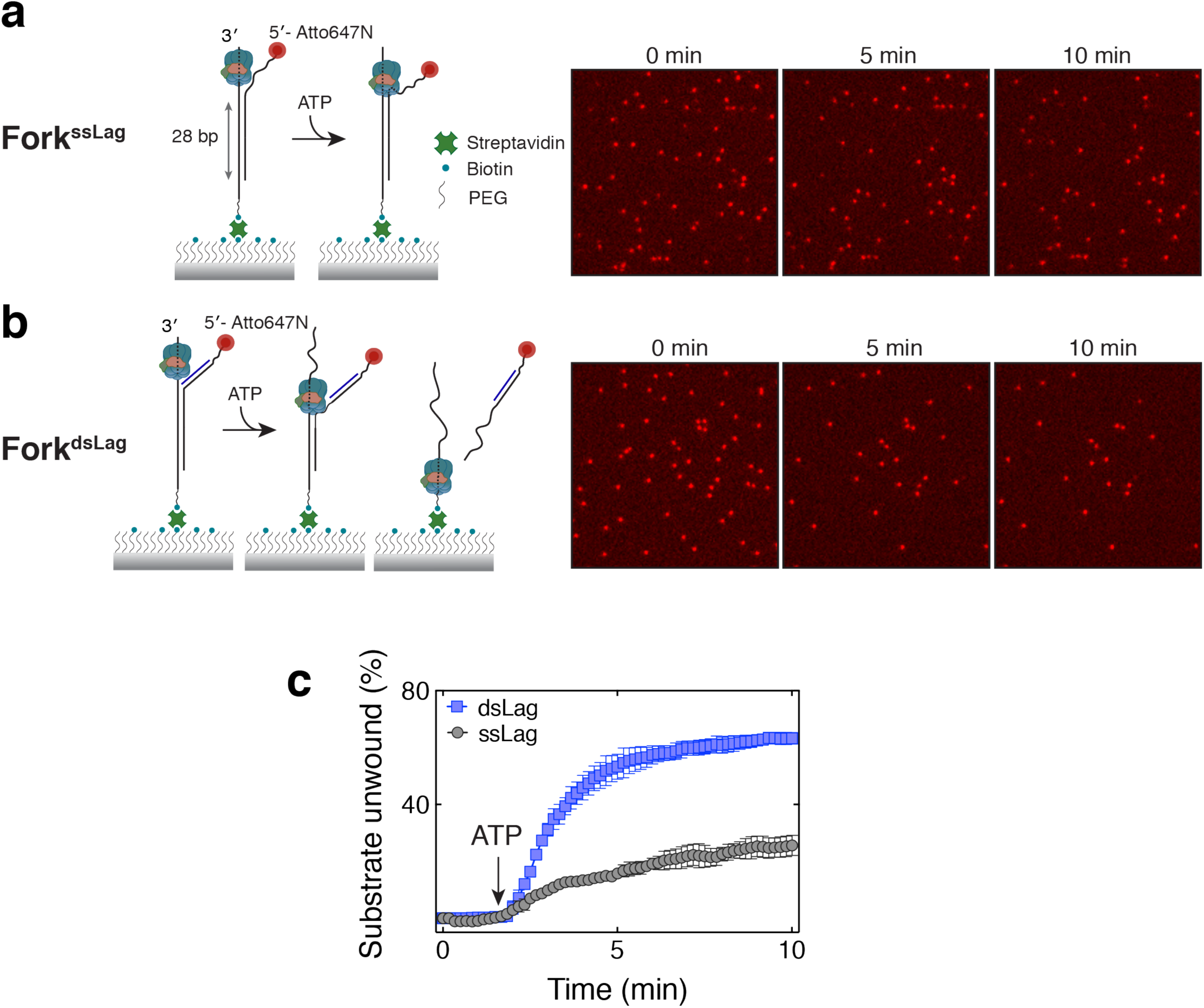
Single-molecule analysis of CMG-catalyzed fork unwinding. **a-b,** Visualizing CMG-driven unwinding of individual Atto-647N-labelled surface-immobilized fork substrates with TIRF microscopy. Unwinding of 28-bp duplex region by CMG leads to dissociation of the fluorescent strand from the surface. Representative fields of view are shown at three time points on Fork^ssLag^ (**a**) and Fork^dsLag^ (**b**) following ATP addition. **c,** Percentage of molecules unwound as a function of time for Fork^ssLag^ (grey, N=1215 molecules analyzed) and Fork^dsLag^ (blue, N=1611 molecules analyzed). Data represent mean ± SD from three independent experiments for each substrate.

### Binding of RPA to the excluded strand promotes DNA unwinding by CMG

At the eukaryotic replication fork, while the leading strand is proximately synthesized behind CMG, lagging-strand replication proceeds by discontinuous synthesis of Okazaki fragments. While the lagging-strand template near the fork junction remains single stranded during fork progression, it is bound by the single-stranded binding protein, replication protein A (RPA), which may prevent CMG from partially encircling the excluded strand in its central pore. This model could also explain how RPA speeds up dsDNA unwinding by CMG seen in our ensemble (Figure 2) and single-molecule experiments (Figure 1) (Burnham et al., 2019; Kose et al., 2019). We tested whether RPA binding to the excluded strand can stimulate CMG’s ability to unwind fork DNA using the fluorescence-quencher labelled DNA unwinding assay. In this experiment, after binding CMG on fork DNA in the presence of ATPγS, RPA was added to bind the 5ʹ tail of the fork prior to addition of ATP. A capture oligonucleotide was also included together with ATP to sequester both excess CMG and RPA binding to DNA once unwinding began. Addition of 25 nM RPA led to 3-fold higher fluorescence signal on Fork^ssLag^ (Figure 5a). At this RPA concentration unwinding was strictly dependent on the presence of CMG (Supplementary Figure 5a). Critically, RPA did not increase unwinding of Fork^dsLag^ by CMG (Figure 5b), corroborating that RPA binding to the lagging-strand arm of Fork^ssLag^ was responsible for the observed stimulation. These results suggest the presence of RPA on the lagging-strand template enhances CMG helicase activity by hindering engagement of the helicase with parental duplex at the fork junction providing mechanistic insight into how RPA speeds up CMG-driven DNA unwinding.

**Figure 5.**
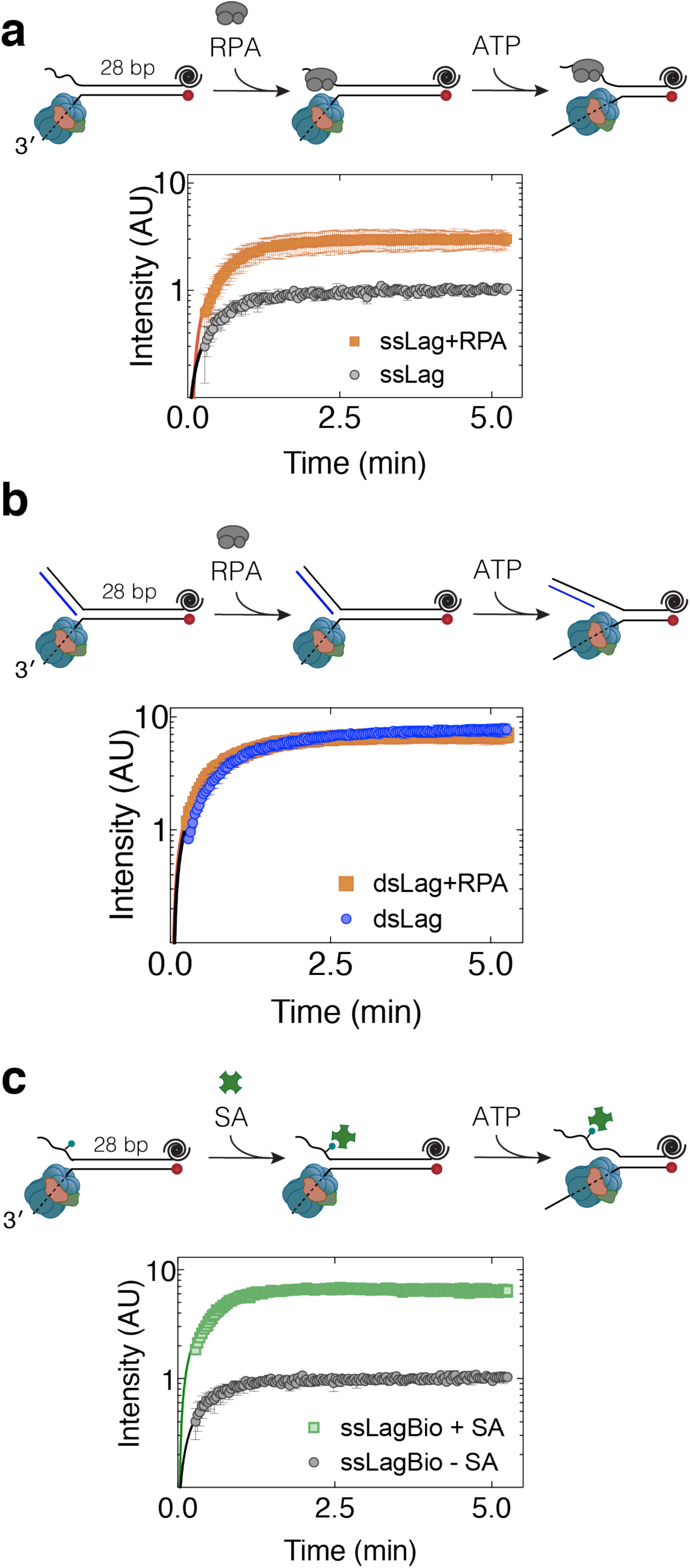
Binding of a protein to the excluded strand promotes DNA unwinding by CMG. **a,** Single turn-over time-course unwinding of Fork^ssLag^ by CMG in the absence (black) and presence (orange) of RPA. **b,** Single turn-over time-course unwinding of Fork^dsLag^ by CMG in the absence (blue) and presence (orange) of RPA. In **a** and **b**, RPA was added after CMG binding to the fork. A competitor oligonucleotide was included with ATP addition to prevent further RPA and CMG binding during unwinding. Data represent mean ± SD (n=3). **c,** CMG-catalyzed single turn-over unwinding of fork DNA containing a single biotin on the lagging-strand arm in the absence (black) and presence (green) of streptavidin (SA). Data represent mean ± SD (n=3). Solid lines are fits to Equation 2 (Methods).

### Streptavidin binding to the excluded strand is sufficient to stimulate CMG helicase activity

Our findings strongly suggest that, when single-stranded, the excluded strand near the fork nexus is inhibitory to fork unwinding by CMG. While one possible reason for the observed inhibition is the partial entrapment of the displaced strand (hence the parental duplex) in the helicase central pore, the other consideration is the interaction of the displaced strand with the outer surface of CMG. If a physical contact between the exterior of CMG and the lagging-strand template adversely affects helicase activity, as suggested for DnaB (Carney et al., 2017), hybridization of a complementary oligonucleotide or binding of RPA to the lagging-strand arm of a fork substrate may stimulate DNA unwinding by disrupting this interaction. To determine whether the inhibition of CMG helicase activity stems from its interaction with the displaced strand through the helicase exterior residues or interior channel, we introduced a biotin-streptavidin complex on the lagging-strand arm of a fork substrate. The streptavidin was bound to biotin on a thymidine base through a 6-carbon spacer, thus is unlikely to interfere between the interaction of the lagging-strand arm and the external surface of the CMG. However, the large size of streptavidin bound to the lagging-strand template is expected to prevent the helicase ring from encircling this strand. Therefore, streptavidin should stimulate fork unwinding only if the observed inhibition was due to CMG encircling the displaced strand at the fork junction and engaging with the parental duplex. In agreement with this model, we found that addition of streptavidin to fork DNA modified with biotin on the 5ʹ tail enhanced CMG helicase activity (Figure 5c) similar to the levels observed with duplexing the 5ʹ tail (Figure 3b). Addition of streptavidin on a non-biotinylated fork substrate had no effect (Supplementary Figures 5b and 5c) indicating that streptavidin promoted CMG-driven fork unwinding by binding to the 5ʹ tail of the fork template. Together, our data suggest that CMG pauses when engaged with the lagging-strand template via its central pore and that this non-productive helicase-fork arrangement is averted by binding of a protein to the lagging-strand template.

### DNA reannealing can free CMG from the duplex-engaged state

It is conceivable that CMG occasionally stalls at the replication fork upon sliding onto duplex DNA when the lagging-strand template adjacent to the replication fork is not occupied by a protein such as RPA. To resume DNA unwinding, the helicase has to exit this paused state. As CMG exhibits a biased random walk during DNA unwinding (Burnham et al., 2019), we reasoned that the helicase should be able to move backwards and revert to a strand exclusion mode if paused due to surrounding a short tract of parental DNA. To investigate whether CMG can exit a duplex-engaged mode, we allowed the helicase to first enter into the trapped state on the fork substrate used in Figure 3 (Fork^ssLag^) consisting of 28-bp dsDNA. To this end, CMG was first bound to Fork^ssLag^ with ATPγS and its translocation on DNA subsequently triggered with ATP. When Comp^Lag^ was added to hybridize to the 5ʹ tail of the fork before (Figure 6a, Comp^Lag^ → ATP, blue) or together with ATP (Supplementary Figure 3e), DNA unwinding was markedly increased compared to the reaction lacking Comp^Lag^. Surprisingly, when introduced 2 minutes after ATP, Comp^Lag^ did not stimulate DNA unwinding (Figure 6a, ATP → Comp^Lag^, red) suggesting that once CMG engages with duplex DNA at the fork nexus, it is incapable of retracting to eject the displaced strand and the parental dsDNA completely from the interior channel. Since the reverse motion of CMG was inferred from DNA rezipping in magnetic-tweezers measurements (Burnham et al., 2019), we envisaged that fork reannealing ahead of CMG may be required to push the helicase backwards to exit a paused state. In this scenario, when using the synthetic fork substrate shown in Figure 6a, CMG would not move in the reverse direction when trapped at the fork junction because this fork substrate contains two non-complementary ssDNA tails, unlike a genuine replication fork. To investigate whether CMG can exit a duplex-engaged mode due to DNA rewinding, we measured its helicase activity on a fork substrate containing 60-bp dsDNA. The 5ʹ tail of this fork substrate consisted of d(GGCA)_10_ that prevents the helicase entering the paused state near the ss-dsDNA junction (Supplementary Figure 3a). The presence of relatively long dsDNA on this substrate makes it likely that CMG will enter the aforementioned stalled state while unwinding the 60-bp duplex section. However, the helicase may escape from the paused state by translocating backwards as a consequence of the two strands in front of the helicase rezipping (Figure 6b). When included before ATP, an oligonucleotide complementary to the excluded strand within the duplex region enhanced unwinding by CMG (Figure 6b, Comp^Lag^ → ATP, blue) suggesting that the helicase engages with dsDNA within the 60-bp parental duplex and stalls during unwinding. Strikingly, CMG-mediated unwinding of this substrate quickly recovered even when Comp^Lag^ was introduced 20 minutes after ATP (Figure 6b, ATP → Comp^Lag^, red). This result is consistent with the notion that if the parental duplex invades into the CMG central channel, reannealing of DNA ahead of the helicase can liberate CMG from this inactive fork-binding mode by promoting backwards translocation.

**Figure 6.**
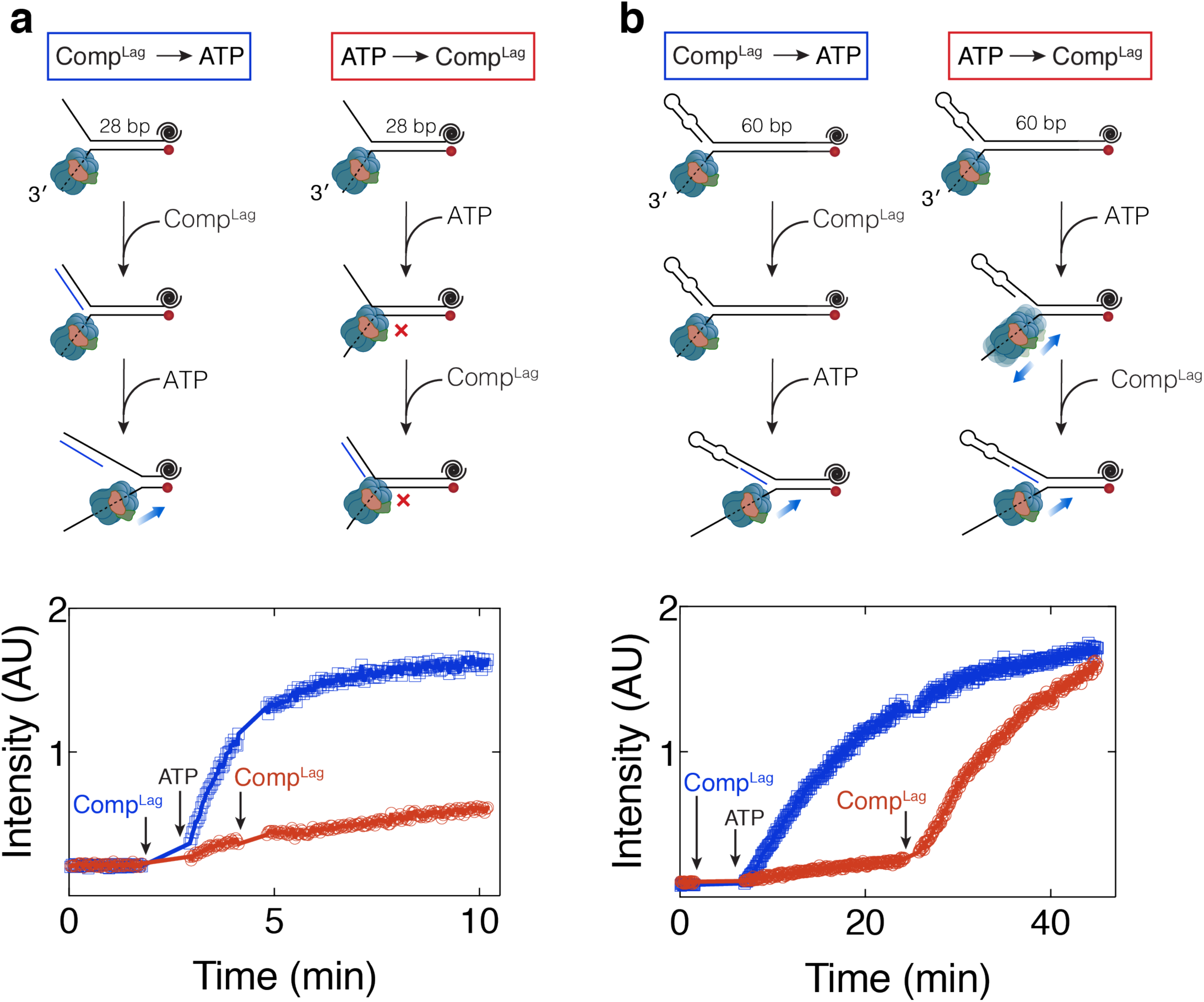
DNA reannealing can free CMG from the duplex-engaged state. **a,** Single turn-over unwinding of fork DNA with a single stranded lagging-strand arm and 28 bp parental duplex. CMG was bound to the fork in the presence of ATPγS. Subsequently ATP was added and unwinding was monitored through Cy5 fluorescence. An oligonucleotide complementary to the lagging-strand arm (Comp^Lag^) was added either before (blue, Comp^Lag^→ATP) or 2 minutes after (red, ATP→Comp^Lag^) ATP. **b,** Single turn-over unwinding of fork DNA with d(GGCA)_10_ lagging-strand arm and 60-bp parental duplex. CMG was bound to the fork in the presence of ATPγS. Subsequently ATP was added and unwinding was monitored through Cy5 fluorescence. An oligonucleotide complementary to the lagging-strand template (Comp^Lag^) within the 60-bp parental dsDNA was added either before (blue, Comp^Lag^→ATP) or 20 minutes after (red, ATP→Comp^Lag^) ATP.

### CMG bypasses a protein directly crosslinked to the excluded strand with no detectable stalling

A recent cryo-EM structure of yeast CMG, where translocation on a fork DNA substrate is blocked by biotin-streptavidin barriers, exhibited a short stretch of dsDNA enclosed by the N-terminal zinc finger (ZF) protrusions of Mcm2-7 (Georgescu et al., 2017). In a similar assay, we showed that the N-terminal end of MCM in *Drosophila* CMG interacts with parental DNA at the fork in an analogous manner (Eickhoff et al., 2019) suggesting that the mode of helicase engagement with DNA is conserved between *Drosophila* and yeast. A possible exit route for the displaced lagging-strand template was proposed to be formed by cavities between MCM ZF protrusions (Eickhoff et al., 2019, Li and O’Donnell, 2018). If the displaced strand exits through a narrow opening between MCM ZF domains during unwinding, CMG is expected to pause when colliding with a bulky obstacle on this strand. We previously demonstrated that while a methyltransferase crosslinked to the lagging-strand template slowed down DNA unwinding by CMG, a covalent lagging-strand streptavidin block did not alter CMG dynamics (Kose et al., 2019). There are two major differences between these two types of DNA-protein crosslinks (DPCs). The first is that methyltransferase interacts with both DNA strands and increases the stability of duplex DNA, hence impeding CMG translocation (Kose et al., 2019). Secondly, while methyltransferase was directly crosslinked to a base, streptavidin was attached to DNA through a flexible linker. Thus, the absence of CMG slowing down at a lagging-strand streptavidin barrier may be due to the linker escaping through the narrow channel between MCM ZF domains. To test this possibility, we used 5-formylcytosine (5fC) modification to form a DPC lacking a linker between DNA and streptavidin (Ji et al., 2017; Li et al., 2017) (Figure 7a). We generated a fork substrate containing a single 5fC-streptavidin crosslink on the lagging-strand template in the middle of 60-bp parental duplex (Supplementary Figure 6). We measured unwinding dynamics using the fluorescence-based single turn-over kinetic unwinding assay as described in Figure 3. Importantly, we found that fork DNA containing a 5fC-streptavidin crosslink was unwound by CMG as efficiently and as rapidly as the non-adducted substrate (Figure 7b). This result suggests that the lagging-strand template is unlikely to be expelled through a tight gap in the N-terminal region of Mcm2-7 during unwinding. Together with our data demonstrating that duplex engagement by CMG severely slows it down during DNA unwinding, we favour a model in which CMG engaged with the fork by encircling parental duplex through MCM ZF domains reflects a stalled helicase. Accordingly, multiple different conformations of parental duplex with respect to the N-terminal region of MCM were observed in *Drosophila* CMG/fork DNA structure suggesting plasticity in duplex engagement by the helicase (Eickhoff et al., 2019).

**Figure 7.**
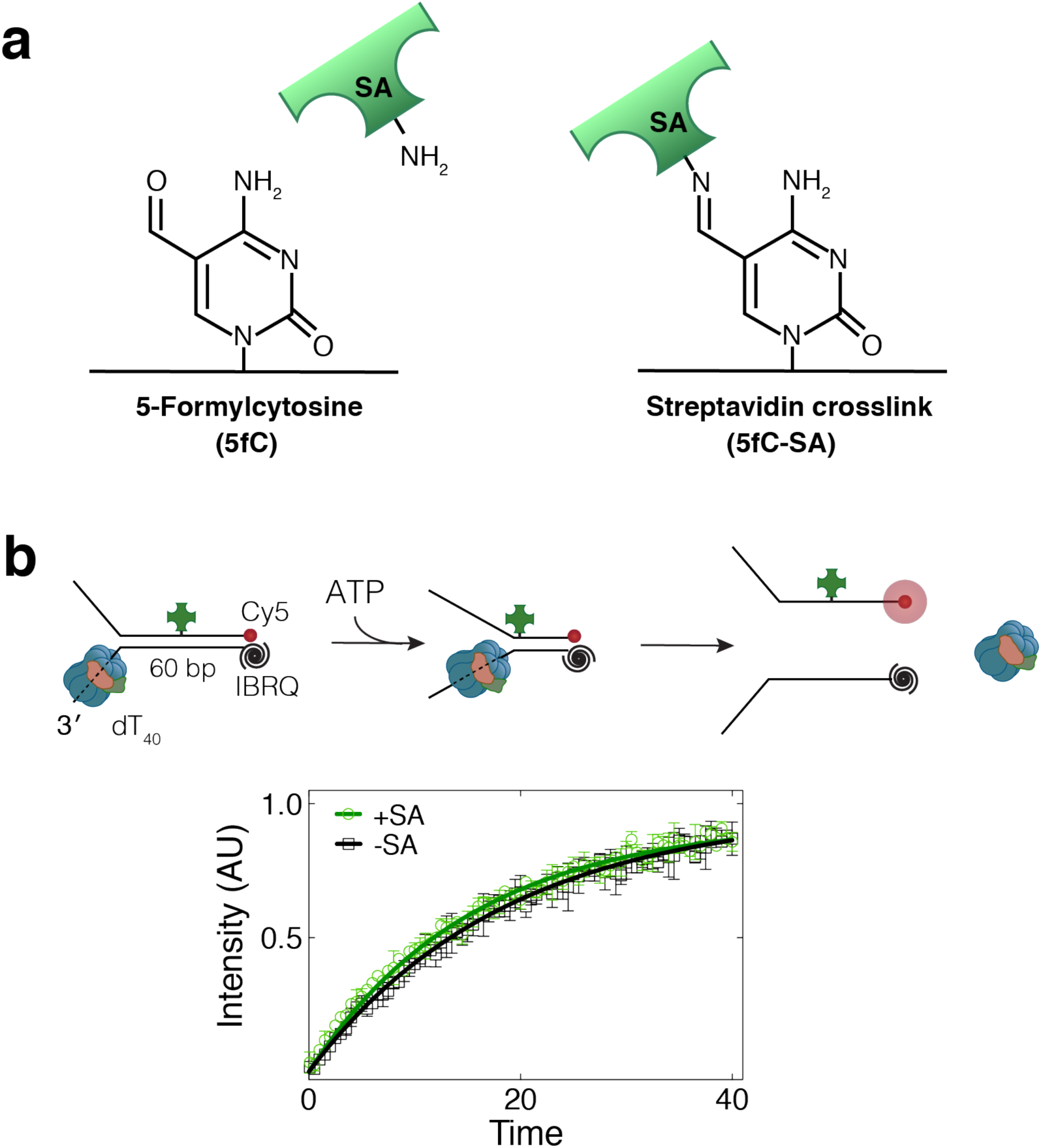
CMG bypasses a protein directly crosslinked to the excluded strand with no detectable stalling. **a,** 5-formyl-cytosine-modified oligonucleotide covalently crosslinks to streptavidin through primary amines. **b,** Single turn-over CMG-mediated unwinding of fork DNA substrates with and without a 5fC-streptavidin crosslink. DNA substrates contained 60-bp duplex region and were modified with Cy5 at the 3ʹ end of the lagging-strand template and Iowa Black RQ (IBRQ) dark quencher at the 5ʹ end of the leading-strand template. Separation of the complementary strands by CMG leads to fluorescence increase, a proxy for DNA unwinding. CMG unwinds 5fC-streptavidin-crosslinked and non-crosslinked substrates with the same kinetics. Data represent mean ± SD (n=3). Solid lines represent fits to Equation 2 (Methods).

## Discussion

Separation of the DNA double helix is catalyzed by the CMG helicase during eukaryotic replication. While isolated CMG complex can unwind short stretches of duplex DNA (tens of bp) and travel long distances on ssDNA, it shows slow or no substantial helicase activity on long dsDNA (thousands of bp) in the absence of additional replication factors (Burnham et al., 2019; Lewis et al., 2017; Wasserman et al., 2019). To discover the origin of poor helicase activity of isolated CMG, we investigated how DNA unwinding by the helicase is regulated by its interaction with the replication fork. We found that interaction of CMG with ss-dsDNA fork junction leads to helicase stalling likely due to partial entrapment of parental duplex within the helicase central channel, which explains the extremely slow translocation of CMG while unwinding the fork.

Impediment of fork unwinding appears to be a common feature of replicative helicases, especially when their central channels are wide enough to encircle dsDNA. Similar to CMG, duplex unwinding by replicative helicases T7 gp4 and *E.coli* DnaB is hindered by their interactions with duplex DNA at the fork junction (Atkinson et al., 2010; Jeong et al., 2013; Kaplan, 2000). Measurement of DNA hairpin unwinding by T4 gp41, T7 gp4 and *E.coli* DnaB with single-molecule magnetic and optical tweezers showed that these replicative helicases unwind dsDNA 5-10-fold slower compared to their ssDNA translocation rates (Johnson et al., 2007; Lionnet et al., 2007; Ribeck et al., 2010). Similarly, while single CMG complexes unwind dsDNA at rates of 0.1-0.5 bps^-1^ (Burnham et al., 2019), they translocate on ssDNA with an average rate of 5-10 nts^-1^ (Wasserman et al., 2019), more than 10-fold variance. In contrast, large T antigen, which has a relatively small central channel, unwinds dsDNA only 2-fold slower than it translocates on ssDNA (Klaue, 2012). These results support a model whereby inhibition of DNA unwinding due to parental duplex engagement is a common feature of replicative helicases and suggests that the likelihood of helicase slowing down at the fork correlates with their central pore size.

A recent structure of the T7 replisome demonstrated that coupling of the gp4 helicase with the leading-strand polymerase positions the helicase central channel axis perpendicular to the parental DNA (Gao et al., 2019). This observation is in line with the prediction that the leading-strand polymerase stimulates DNA unwinding by gp4 by obstructing helicase engagement with duplex DNA. We speculate that faster fork rates seen in DnaB when coupled to the polymerase (Kim et al., 1996) may be through the same mechanism. Unlike bacteriophage and bacteria, the leading-strand polymerase, polymerase epsilon (Pol ε), is placed behind the helicase in eukaryotes. Thus, leading-strand synthesis would not hinder CMG invading onto parental DNA. On the contrary, Pol ε binding to DNA behind CMG may further lead to fork pausing because CMG backtracking appears to be essential to rescue the helicase from duplexed-engaged conformation (Figure 6), which may explain CMG pausing at protein barriers in a Pol ε-dependent manner (Hizume et al., 2018).

In this work, we demonstrate that the presence of RPA on the lagging-strand arm of a fork substrate prevents helicase engagement with the parental duplex near the fork junction. Therefore, we propose that binding of RPA to the lagging-strand template is essential for proper replication fork progression. Accordingly, we show that single CMG complexes can processively unwind thousands of base pairs of DNA in the presence of RPA at a rate similar to the speed of CMG on ssDNA (Wasserman et al., 2019). Because RPA can diffuse along ssDNA (Kemmerich et al., 2016), we envisage that new RPA from solution does not need to immediately bind to the emerging lagging-strand template near the helicase to support unwinding. Furthermore, like RPA, other replication factors such as polymerase alpha-primase, Mcm10, and AND-1 (Ctf4), may also promote DNA unwinding by keeping the helicase from interacting with the parental duplex. The inability of CMG to drive extensive DNA unwinding in the absence of RPA should be beneficial for cells. Specifically, under conditions of RPA exhaustion, capture of the parental duplex by the N-terminal region of the MCM pore would lead to helicase slowing and prevent accumulation of unprotected ssDNA. Upon recovery of RPA, we propose that re-annealing of the parental duplex would drive CMG backwards and restore DNA unwinding (Figure 8). The plasticity of the eukaryotic replicative helicase to move backwards may be particularly important to keep the replisome in place during replication fork reversal (Neelsen and Lopes, 2015) such that replication fork restart can take place without the need for new helicase loading onto DNA, which is inhibited in S phase (Arias and Walter, 2007). Additionally, 5’-to-3’ helicases may also help to rescue a duplex-engaged CMG by acting on the excluded strand at the replication fork.

**Figure 8.**
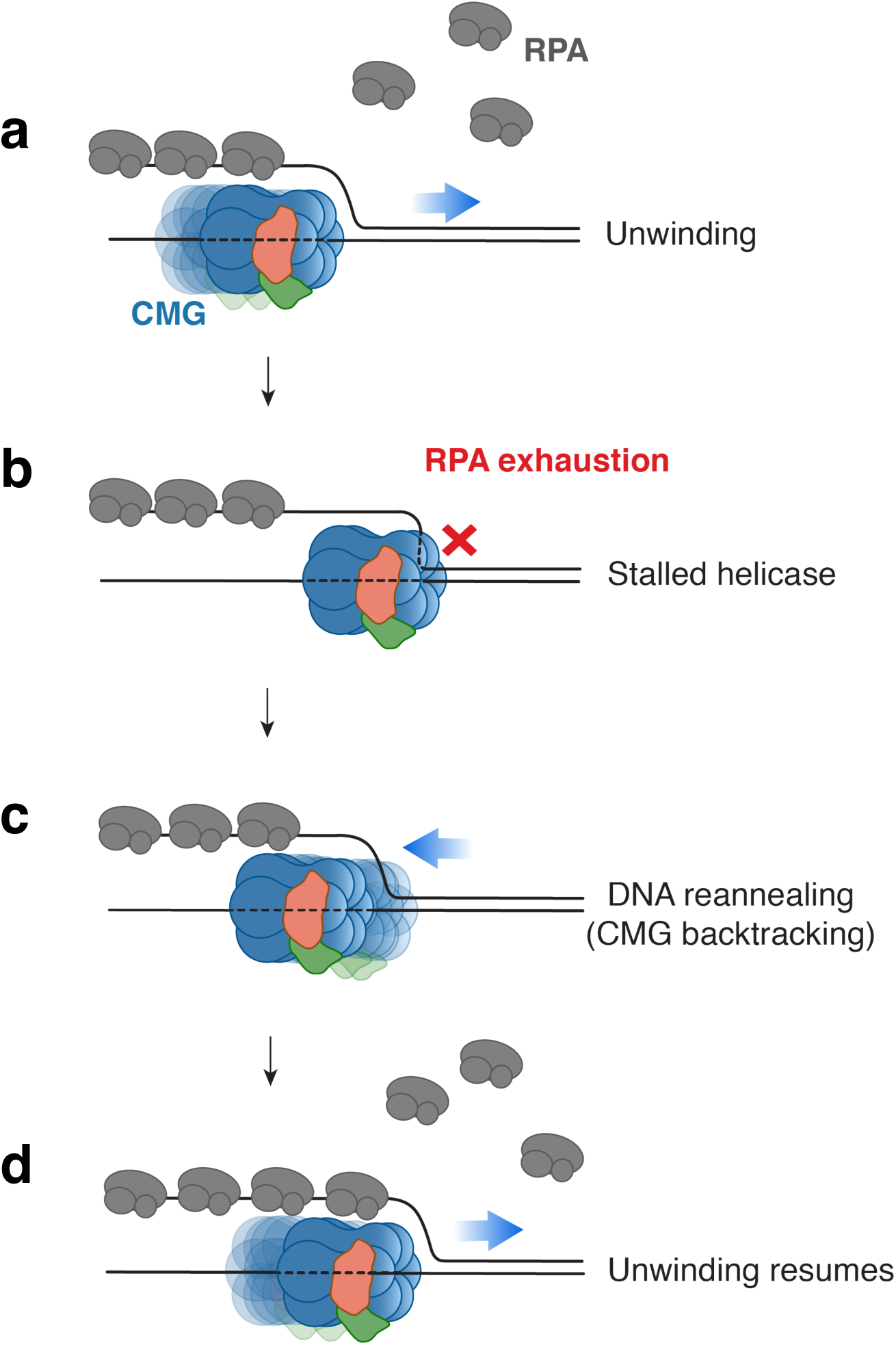
Model for RPA-facilitated DNA unwinding by CMG. **a,** Binding of RPA to the lagging-strand template prevents CMG from engaging with duplex DNA at the fork junction. Thus, in the presence of RPA, CMG unwinds dsDNA at rates similar to its ssDNA translocation rate. **b,** When RPA is not available, duplex DNA engagement leads to helicase stalling/slowing down. **c,** DNA rezipping-induced CMG backtracking can rescue a stalled helicase. **d,** Faster unwinding rates are restored upon availability of RPA.

It is not clear whether the recent CMG structures in complex with fork DNA exhibiting dsDNA within the MCM ZF domains represent the helicase in an active fork unwinding state or a paused state (Eickhoff et al., 2019; Georgescu et al., 2017). Because CMG can unwind dsDNA much more efficiently with the help of RPA, a high-resolution structure of CMG while unwinding duplex DNA in the presence of RPA should further elucidate how the helicase engages with DNA in the eukaryotic replisome.

Rate of DNA unwinding by *Drosophila* CMG in the presence of RPA (4.5 bps^-1^, Figure 1) is still an order of magnitude lower than average cellular replication fork rates in *Drosophila melanogaster* (∼43 bps^-1^) (Blumenthal et al., 1974). The observation that unwinding rates are reduced when CMG uncouples from the leading-strand synthesis (Sparks et al., 2019; Taylor and Yeeles, 2019) indicates that pol ε increases replication fork rates. In yeast, Mrc1-Tof1-Csm3 complex (Claspin-Timeless-Tipin in metazoans) is reported to further enhance fork speed (Lewis et al., 2017; Yeeles et al., 2017). The molecular mechanism by which pol ε and Mrc1-Tof1-Csm3 speed up fork progression is unclear. Using the single-molecule DNA unwinding assays described here, it will be important to test whether these factors can stimulate DNA unwinding by CMG in the absence of DNA synthesis.

## Methods

### DNA substrates

Supplementary Table 1 includes a list of oligonucleotides (oligos) used to prepare various DNA substrates and the corresponding figure numbers. The oligonucleotide sequences can be found in Supplementary Table 2.

To prepare fork DNA substrates, oligos (10 µM final), as listed in Supplementary Tables 1 and 2, were mixed in STE buffer (10 mM Tris-HCl pH 8.0, 100 mM NaCl, 1 mM EDTA), heated to 85°C, and allowed to slowly cool down to room temperature. DNA substrates were ligated with T4 DNA Ligase (NEB) to seal nicks when necessary. The resulting products were separated on 8% polyacrylamide gel electrophoresis (PAGE) and purified by electroelution, unless stated otherwise. Electroelution was performed using 3.5 kDa MWCO dialysis tubing (Spectra/Por, Spectrum Labs) in 1× TBE buffer.

Fork DNA containing 236-bp duplex and internal Cy5 (Figure 2a) was made by PCR amplification of 314-bp fragment from pHY10 (Duxin et al., 2014) using primers Oligo-2 and Oligo-3 in the presence of Cy5-dCTP (GE Healthcare). The PCR reaction was purified using QIAquick PCR purification kit (Qiagen), digested with NcoI, and separated on 2% agarose gel. The resulting 211-bp fragment with 5’-CATG-3’ overhang was purified using gel extraction kit (Qiagen), ligated to 10-fold excess of short fork generated by annealing Oligo-4 and Oligo-5. The final product was separated on 8% PAGE and purified via electroelution. To make 252-bp duplex fork with downstream 28-bp duplex and 3’-Cy5-modified strand (Figure 2b), 314-bp fragment from pHY10 was PCR amplified using primers Oligo-2 and Oligo-17. After purifying the reaction with QIAquick PCR purification kit, the substrate was cut with NcoI and HindIII (NEB), purified after separating on 2% agarose, and ligated to oligos that were annealed and purified from 8% PAGE. While NcoI-digested end was ligated to Oligo-4/Oligo-5, HindIII-treated end was ligated to Oligo-15/Oligo-16/Oligo-Cy5-6. The final reaction was separated on 2.5% agarose and purified using electroelution. The substrate lacking the 40-nt 3’ polyT tail (Supplementary Figure 2) was made using exactly the same protocol except Oligo-5 was replaced with Oligo-18.

To prepare 5fC-streptavidin-crosslinked fork substrate (Figure 7), 1 nmol 5fC-modified oligo (Oligo-5fC) was incubated with 1 mg streptavidin (Sigma) in 100 µl PBS at 55°C for 4 hours followed by 37°C for 12 hours. DNA was separated on 8% PAGE in 1x TBE. The band corresponding to streptavidin-crosslinked oligo (1-2% of the reaction) was excised, purified via electroelution, annealed and ligated to other oligos as listed in Supplementary Table 1. The streptavidin-crosslinked fork was purified again by electroelution after separating on 8% PAGE.

10-kb linear DNA used in single-molecule assays (Figure 1) was generated by treating λ DNA with ApaI (NEB). 10-kb fragment from ApaI-cut λ DNA was purified from a 0.5% agarose gel using Monarch gel extraction kit (NEB). We generated the fork end containing a 5ʹ biotin and a Cy3 on the 3ʹ dT_40_ tail by annealing Oligo-Bio-4 and Oligo-Cy3-1, which leaves a 12-nt ssDNA complementary to one end of the 10 kb λ-ApaI fragment. To label the other end of 10 kb DNA with digoxigenin, a 0.5 kb fragment was PCR amplified from pUC19 vector with primers Oligo-1 and Oligo-8 in the presence of digoxigenin-11-dUTP (Roche). Digoxigenin-modified PCR substrate was then digested with ApaI and purified using QIAquick PCR purification kit. 2-3 µg of 10 kb DNA was mixed with 10-fold molar excess of Cy3-biotin-labelled fork substrate and digoxigenin-modified PCR template, and ligated with T4 DNA ligase. The reaction was mixed with 15 µg streptavidin for attachment to the biotin end of the fork, separated on 0.5% agarose in 1xTBE, and purified via electroeleution.

### Protein expression and purification

#### Drosophila melanogaster CMG

*Drosophila* CMG was expressed, and purified as described in (Ilves et al., 2010; Kose et al., 2019). Each pFastBac1 (pFB) vector containing a single subunit of *Drosophila* CMG was transformed into DH10Bac *E.coli* competent cells (Thermo Fisher) to generate bacmids. Mcm3 subunit contained an N-terminal Flag tag for purification. Sf21 cells (10^6^/ml) were used for the initial transfection (P1 stage), and in the subsequent virus amplification stage to make P2 stocks using Sf-900TM III SFM insect cell medium (Invitrogen/Gibco). In all virus amplification stages, cells were incubated at 27°C while shaking at 120 rpm. To further amplify virus stocks (P3 stage), 100 ml Sf9 cell cultures (0.5×10^5^/ml) in Graces medium supplemented with 10% FCS for each subunit were infected with 0.5 ml P2 viruses and incubated in 500 ml Erlenmeyer sterile flasks (Corning) for ∼100 hours. Total of 200 ml P3 viruses for each subunit were used to infect 4 L of Hi5 cells (10^6^/ml) with a multiplicity of infection (MOI) of 5. Hi5 cells were divided into 500 ml aliquots using sterile 2 L roller bottles (Corning). After 48 hours, cells were harvested by centrifuging at 4,500 x g. Cell pellets were first washed with phosphate-buffered saline (PBS) supplemented with 5 mM MgCl_2_, resuspended in 200 ml Buffer C-Res (25 mM HEPES pH 7.5, 1 mM EDTA, 1 mM EGTA, 0.02% Tween-20, 10% glycerol, 15 mM KCl, 2 mM MgCl_2_, 2 mM β-ME (2-mercaptoethanol), PI tablets) (50 ml buffer per 1 L Hi5 cell culture), and frozen in 10 ml aliquots on dry ice. Cell pellets were stored in −80°C.

All purification steps were performed at 4°C unless specified otherwise. Frozen cell pellets were thawed in lukewarm water, and lysed by 60-70 strokes using cell homogenizer (Wheaton, 40 ml Dounce Tissue Grinder) on ice. Cell debris was removed by centrifugation at 24,000 × g for 10 minutes. The supernatant was collected, and incubated with M2 agarose Flag beads (Sigma Aldrich) equilibrated with Buffer C (25 mM HEPES pH 7.5, 1 mM EDTA, 1 mM EGTA, 0.02% Tween-20, 10% glycerol, 1 mM DTT) for 2.5 hours. CMG was eluted from beads by incubating with Buffer C-100 (25 mM HEPES pH 7.5, 1 mM EDTA, 1 mM EGTA, 0.02% Tween-20, 10% glycerol, 100 mM KCl, 1 mM DTT) supplemented with 200 µg/ml peptide (DYKDDDDK) for 15 minutes at room temperature. The eluate was passed through 1 ml HiTrap SPFF column (GE Healthcare) equilibrated with Buffer C-100. CMG was separated with 100-550 mM KCl gradient using 5/50GL MonoQ column (GE Healthcare). Fractions containing CMG were pooled, diluted to 150 mM KCl, and loaded onto MonoQ PC 1.6/5GL (GE Healthcare) column equilibrated with Buffer C-150 (25 mM HEPES pH 7.5, 1 mM EDTA, 1 mM EGTA, 10% glycerol, 150 mM KCl, 1 mM DTT) to concentrate the sample. 150-550 mM KCl gradient was applied to elute CMG. Fractions containing CMG were pooled, and dialyzed against CMG-dialysis buffer (25 mM HEPES pH 7.5, 50 mM sodium acetate, 10 mM magnesium acetate, 10% glycerol, 1 mM DTT) for 2 hours.

To prepare the fluorescently-labelled CMG construct, site-directed mutagenesis (QuikChange, Agilent) was applied to pFB-Mcm3 vector to insert a TEV cleavage site (ENLYFQG) followed by 4 Gly residues downstream of the N-terminal Flag tag on Mcm3 for subsequent Sortase-mediated labelling. The construct was expressed and purified as before with slight modifications. After collected from 5/50GL MonoQ column, 1 ml of the sample was mixed with 50 μl TEV protease (1 mg/ml, EZCut TEV Protease, Biovision), and the mixture was dialyzed against Buffer C-100 overnight at 4°C. TEV-treated sample was supplemented with 50 μM peptide NH_2_-CHHHHHHHHHHLPETGG-COOH, labelled with LD-655-MAL (Lumidyne Technologies) on the cysteine residue, 10 μg/ml Sortase enzyme and 5 mM CaCl_2_, and incubated 30 minutes at 4°C. Free peptide was removed by separating the sample through gel filtration (Superdex 200 Increase 10/300 GL) in Buffer C-100. The labelled construct was concentrated on MonoQ PC 1.6/5GL and dialyzed as described above.

#### Sortase

pET29-based expression vector for C-terminally 6xHis-tagged Sortase A pentamutant (Sortase P94R/D160N/D165A/K190E/K196T) (Chen et al., 2011) was obtained from Dr. David Liu (Harvard University). *E.coli* BL21(DE3) competent cells transformed with the expression vector were grown at 37°C in LB with 50 μg/ml kanamycin until OD=0.5-0.8 and induced with 0.4 mM isopropyl β-D-1-thiogalactopyranoside (IPTG) for 3 hours at 30°C. Cells were harvested by centrifugation and resuspended in Sortase-lysis buffer (50 mM Tris pH 8.0, 300 mM NaCl, 1 mM MgCl_2_, 2 units/ml DNAseI (NEB), 260 nM aprotinin, 1.2 μM leupeptin, 1 mM PMSF). After lysing cells by sonication, cells were centrifuged at 7,500 × g for 20 minutes. The supernatant was supplemented with 10 mM imidazole, mixed with Ni-NTA agarose (1 ml bed volume pre-washed with Sortase-lysis buffer per 8 ml lysate), and incubated for 1 hour on a rotary shaker. The lysate-Ni-NTA mixture was loaded into a gravity column and washed twice with 5 bed volumes of Sortase-wash buffer (50 mM Tris pH 8.0, 300 mM NaCl, 1 mM MgCl_2_, 20 mM imidazole). The protein was eluted from Ni-NTA beads four rounds each with one bed volume of Sortase-elution buffer (50 mM Tris pH 8.0, 300 mM NaCl, 1 mM MgCl_2_, 250 mM imidazole). Fractions containing Sortase were combined and dialyzed against Sortase-storage buffer (25 mM Tris pH 7.0, 150 mM NaCl, 10% glycerol).

#### Large T antigen

Recombinant SV40 large T antigen was made using the baculovirus expression system and purified using a monoclonal antibody as described previously (Yardimci et al., 2012). pFB with the full length large T-antigen was used to make the baculovirus. Sf21 cells maintained in SF-900-III were infected with the virus (2 × 10^8^ pfu/ml) using an MOI of 0.1 for 72 hours. Cell pellets were re-suspended in 10 pellet volumes of L-Tag-resuspension buffer (20 mM Tris pH 9.0, 300 mM NaCl, 1 mM EDTA, 10% glycerol, 0.5% NP-40, 0.1 mM DTT) and incubated on ice for 15 minutes. The suspension was centrifuged at 25,000 × g for 15 minutes. 0.5 volume of L-Tag-neutralisation buffer (100 mM Tris pH 6.8, 300 mM NaCl, 1 mM EDTA, 10% glycerol, 0.5% NP-40, 0.1 mM DTT) was added and mixed.

The protein was affinity purified using a mouse monoclonal antibody (PAb419). Antibody was coupled to Protein A sepharose beads (5 mg/ml) in PBS, incubated rotating overnight at 4°C. Beads were washed with 15 ml 0.1M sodium borate buffer, pH 9.0, and re-suspended in 2 ml of the same buffer. To crosslink antibody to beads, 20 ml of 12.5 mg/ml dimethyl pimelimidate (DMP) (Sigma) was mixed with the beads and incubated at room temperature rotating for 1 hour. Coupling reaction was quenched by incubating beads in 0.2 M ethanolamine pH 8.0, rotating for 1 hour at room temperature. Large T antigen suspended in neutralisation buffer was first loaded onto protein A-only column, equilibrated with L-Tag-loading buffer (20 mM Tris pH 8.0, 300 mM NaCl, 1 mM EDTA, 10% glycerol, 0.5% NP-40, 0.1 mM DTT). Flow through was then loaded onto PAb419-conjugated column equilibrated with L-Tag-loading buffer. The column was washed with 50 ml L-Tag-loading buffer, then 50 ml L-Tag-wash buffer (50 mM Tris, pH 8.0, 1M NaCl, 1mM EDTA, 10% glycerol), followed by 20 ml L-Tag-EG buffer (50 mM Tris, pH 8.5, 500mM NaCl, 1mM EDTA, 10% glycerol, 10% ethylene glycol). Large T antigen was eluted with 5–10 ml of L-Tag-elution buffer (50 mM Tris, pH 8.5, 1M NaCl, 10mM MgCl_2_, 1mM EDTA, 10% glycerol, 55% ethylene glycol) and dialyzed overnight into L-Tag-dialysis buffer (20 mM Tris pH 8.0, 10 mM NaCl, 1 mM EDTA, 50% glycerol, 1 mM DTT).

#### NS3 helicase

The expression plasmid containing N-terminal His-SUMO-tagged helicase domain of NS3 (SUMO-NS3h) gene was obtained from Charles Rice (Rockefeller University). NS3h was expressed, and purified as described in (Gu and Rice, 2010). NS3h plasmid was first transformed into Rosetta2 (DE3) competent cells (Novagen). Cells were grown in LB complemented with 50 µg/ml kanamycin and 34 µg/ml chloramphenicol at 37°C. Shaking was interrupted at OD=0.6, and the culture was kept at 4°C for 30 minutes before induction. Cells were induced with 0.4 mM IPTG, transferred to a 16°C incubator and left for 20 hours shaking at 240 rpm. Cells were harvested by centrifugation at 7,500 × g for 10 minutes. Pellets were kept in −80°C.

To purify SUMO-NS3h, cell pellets were resuspended in 100 ml NS3-lysis buffer (20 mM Tris-HCl pH 8.5, 500 mM NaCl, 1 mM β-ME, 10 mM imidazole, 50 µg/ml lysozyme, 2.5 µg/ml RNase A) and incubated on ice for 20 minutes. Next, cells were stirred for 30 minutes at 4°C and subsequently sonicated (3 seconds on, 10 seconds off, 20 cycles). Cell debris was removed by centrifugation at 24,000 × g for 30 minutes. The supernatant was mixed and incubated with 3 ml Ni-NTA agarose beads (Qiagen) for 1 hour. The mixture was transferred to a 20-ml disposable column (Bio-Rad) to collect beads. The column was washed with NS3-wash buffer (20 mM Tris-HCl pH 8.5, 500 mM NaCl, 20 mM imidazole, 1 mM β-ME). SUMO-NS3h was eluted with NS3-Elution buffer (20 mM Tris-HCl pH 8.5, 150 mM NaCl, 250 mM imidazole, 1 mM β-ME). The eluate was loaded onto 5ml HiTrap Q HP column (GE Healthcare) equilibrated with NS3 Buffer A (20 mM Tris-HCl pH 8.0, 100 mM NaCl, 1 mM β-ME). SUMO-NS3h was separated using a 100-500 mM NaCl gradient. Fractions containing SUMO-NS3h was pooled and concentrated to ∼2.5 mg/ml using 30 kDa MWCO spin concentrator (Vivaspin).

To cleave the SUMO tag, SUMO-NS3h was diluted with NS3 Buffer D (20 mM Tris-HCl pH 8.0, 50 mM NaCl, 20% glycerol, 1 mM β-ME) to obtain 100 mM NaCl. 40 units of His-tagged SUMO protease (Invitrogen) was mixed with 1 mg SUMO-NS3h and incubated for 3 hours at 4°C. The sample was then loaded onto 1 ml HisTrap FF column (GE Healthcare) to remove cleaved SUMO tag and SUMO protease. The flow-through which contained cleaved NS3h was collected and concentrated to 0.25 mg/ml using spin concentrator (Vivaspin, 30 kDa MWCO).

#### Human RPA

The expression plasmids for 6xHis-tagged non-fluorescent and EGFP-fused human RPA were obtained Mauro Modesti (Cancer Research Center of Marseille, CNRS). Expression and purification of both RPA constructs were performed as described in (Modesti, 2011). The plasmid was transformed into Rosetta/pLysS competent cells (Novagen). Cells were grown in LB media supplemented with 100 µg/ml ampicillin and 34 µg/ml chloramphenicol at 37°C. At OD=0.5, the temperature was reduced to 15°C, RPA expression was induced with 1 mM IPTG, cells were further incubated for 20 hours. Cells were then harvested by centrifugation at 3,500 × g for 30 minutes. Supernatant was removed, and the cell pellet was washed with PBS. Pellets were stored in −80°C.

To purify RPA, cell pellets were thawed in lukewarm water and resuspended in RPA-lysis buffer A (40 mM Tris-HCl pH 7.5, 1 M NaCl, 20% glycerol, 4 mM β-ME, 10 mM imidazole) supplemented with EDTA-free PI tablets. The suspended pellet was sonicated (3 seconds on, 10 seconds off, 20 cycles) to brake the cells, and cell debris was removed by centrifugation at 20,000 × g for 1 hour. Supernatant was filtered through 0.45 µm syringe filters (Millipore) and loaded onto 1 ml HisTrap FF (GE Healthcare) column equilibrated with RPA-lysis buffer B (20 mM Tris-HCl pH 7.5, 500 mM NaCl, 2 mM β-ME, 20% glycerol, 10 mM imidazole, 1 mM DTT). RPA was eluted by applying linear gradient of 10-300 mM imidazole. Fractions containing RPA were pooled and dialyzed against RPA-dialysis buffer A (20 mM Tris-HCl pH 7.5, 50 mM KCl, 0.5 mM EDTA, 10% glycerol, 1mM DTT) using 3.5 kDa MWCO dialysis tubing (Generon) overnight. Dialyzed sample was filtered using 0.22 µm syringe filter (Millipore) as precipitation occurred. The sample was loaded onto HiTrap Heparin column (GE Healthcare) equilibrated with RPA-dialysis buffer A. The protein was eluted by applying 50-500 mM KCl gradient. Fractions containing RPA were pooled and dialyzed into RPA-dialysis buffer B (20 mM Tris-HCl pH 7.5, 50 mM KCl, 0.5 mM EDTA, 25% glycerol, 1 mM DTT) using 3.5 kDa MWCO dialysis tubing (Generon) for 2 hours.

### Gel-based DNA unwinding assays

To analyze unwinding of Cy5-labelled fork DNA containing 50 bp duplex region (Supplementary Figure 3a), 3 nM DNA was incubated with 50 nM CMG in CMG-binding buffer (25 mM HEPES pH 7.5, 5 mM NaCl, 10 mM magnesium acetate, 5 mM DTT, 0.1 mg/ml BSA) supplemented with 0.1 mM ATPγS at 37°C for 1-2 hours in a 6 µl total volume. 6 µl of ATP mix (CMG-binding buffer supplemented with 5 mM ATP) was added to initiate unwinding and samples were incubated at 30°C for further 10 minutes. Reactions were terminated by addition of 3 µl stop buffer (0.5% SDS, 20 mM EDTA). To prevent aggregation of CMG-bound DNA that results in some DNA being stuck in the well during electrophoresis, 25 μM of 40-nt poly-T oligo (Oligo-6) was included in the stop buffer. Reactions were separated on 8% PAGE in 1x TBE.

To measure unwinding of internally Cy5-modified 236 bp long DNA substrate (Figure 2a), 5 nM fork substrate was incubated with 20 nM CMG in CMG-binding buffer supplemented with 0.1 mM ATPγS at 37°C for 1-2 hours. ATP and RPA was added to final concentrations of 2.5 mM and 150 nM, respectively. After incubating for 1 hour at 30°C, the reactions were stopped with 0.1% SDS, and separated on 3% agarose in 1× TBE supplemented with 0.1% SDS. DNA unwinding assays with 252-bp duplex fork and downstream Cy5-modified strand (Figure 2b) were performed similarly. When RPA was omitted from ATP-containing buffer (Figure 2c), oligonucleotides Oligo-6 (to capture free CMG) and Oligo-21 (to prevent Cy5 strand reannealing to long DNA) were added each at a final concentration of 250 nM. The reactions were incubated at 30°C for indicated times, quenched by adding 0.1% SDS. The stop solution for reactions containing RPA was also supplemented with Oligo-21 (250 nM final) to prevent re-annealing of Cy5-modified strand upon RPA dissociation. Finally, DNA was separated on 8% PAGE in 1× TBE supplemented with 0.1% SDS. To quantify CMG-dependent unwinding in the presence of RPA, the data was corrected for Cy5-modified strand displaced by RPA alone.

Gels were imaged on Fujifilm, SLA-5000 scanner using 635-nm laser and Fujifilm LPR/R665 filter. Band intensities were quantified using ImageJ.

### Real-time DNA unwinding measurements on plate reader

5 nM of Cy5/quencher dual-labelled fork substrates were incubated with 50 nM CMG in CMG-binding buffer in the presence of 0.1 mM ATPγS at 37°C for 1-2 hours. To avoid non-specific binding of CMG and DNA to microplate wells (Nunc 384 shallow well plate, black, 264705), wells were pre-blocked by incubation with CMG-binding buffer supplemented with 1 mg/ml BSA for at least 30 minutes. 5 µl of CMG-DNA mixture was transferred to a well, and 15 µl of ATP mix (CMG-binding buffer containing 3.3 mM ATP) supplemented with 1.5 µM 40-nt polyT oligo (Oligo-6) was added to the reaction. Unwinding assays performed with large T antigen was essentially the same and contained 0.1 mg/ml large T antigen in DNA/helicase binding reaction. For unwinding assays with NS3h, 5 nM of DNA substrate was mixed with 200 nM of NS3h in NS3-MOPS buffer (20 mM MOPS-NaOH pH 6.5, 30 mM NaCl, 3 mM MgCl_2_, 1% glycerol, 0.2% Triton X-100, 1 mM DTT), incubated for 1 hour at 37°C. 5 µl of NS3h/DNA mix was transferred to a well and 15 µl of ATP mix (3.3 mM ATP in NS3-MOPS buffer) supplemented with 1.5 µM 40-nt polyT oligo (Oligo-6) was added start unwinding. When measuring unwinding of fork DNA that contains 28 bp duplex (Figures 3, 5, and 6a), ATP mix was also supplemented with 0.5 µM Oligo-13 to prevent re-annealing of the Cy5-labelled strand to the quencher-modified strand. Comp^Lag^ used for fork substrates containing 28-bp (Figures 3e and 6a) and 60-bp (Figure 6b) duplex regions were Oligo-12 and Oligo-20, respectively.

Cy5 fluorescence intensity was recorded on a PHERAstar FS (BMG Labtech) with excitation and emission wavelengths of 640 and 680 nm, respectively. Data was collected with 2 or 5 seconds intervals with 10 flashes/measurement at 25°C. Measured signals were normalized using Prism 7, and plotted as a function of time.

### Single-molecule DNA unwinding assays

Microfluidic flow cells used in single-molecule assays were made by sandwiching double-sided tape (TESA SE, TESA 4965) between a coverslip, which was coated with PEG and PEG-biotin (Laysan Bio), and a non-functionalized glass slide as described in detail (Yardimci et al., 2012a).

#### Unwinding of Atto647N-labelled short fork substrates

For single-molecule analysis of Atto647N-labelled fork DNA, coverslip surface was first coated with streptavidin by drawing 0.2 mg/ml streptavidin in PBS into the microfluidic flow chamber using a syringe pump (Harvard Apparatus), and incubating for 20 minutes. The flow chamber was extensively washed with blocking buffer (20 mM Tris-HCl pH 7.5, 50 mM NaCl, 2 mM EDTA, 0.2 mg/ml BSA) to remove excess streptavidin. The channel was washed with DNA-dilution buffer (20 mM Tris-HCl pH 7.5, 50 mM NaCl, 2 mM MgCl_2_, 2 mM EDTA, 0.05 mg/ml BSA). Subsequently, 10 pM Atto647N-labelled fork DNA (Fork^ssLag^-TIRF) containing biotin at one end was introduced in DNA-dilution buffer and incubated for 2-3 min for binding to the surface. The flow channel was washed with DNA dilution buffer to remove unbound DNA molecules. 40 nM CMG in 30 µl of TIRF-CMG-loading buffer A (10 mM Tris-HCl pH 7.5, 12 mM MgCl_2_, 15 mM NaCl, 10% glycerol, 0.8 mg/ml BSA, 12.5 mM DTT, 0.3 mM ATPγS) was drawn into the chamber, and incubated for 1 hour. To reduce photobleaching, the flow channel was washed with Atto-imaging buffer (10 mM Tris-HCl pH 7.5, 12 mM MgCl_2_, 15 mM NaCl, 10% glycerol, 0.8 mg/ml BSA, 12.5 mM DTT, 1% glucose, 0.02 mg/ml glucose oxidase (Sigma), 0.04 mg/ml catalase (Sigma)) supplemented with 0.3 mM ATPγS. Surface-tethered DNA was imaged using 647-nm laser at 10-second intervals. After imaging 80 seconds in ATPγS-containing buffer, unwinding was initiated by drawing Atto-imaging buffer containing 3.3 mM ATP into the channel while continuing to collect images. To perform unwinding assays with Fork^dsLag^ substrate, after immobilizing Fork^ssLag^ on the coverslip surface, 50 nM Oligo-12 was introduced in DNA-dilution buffer and incubated for 10 minutes. Flow cell was then washed with DNA-dilution buffer to remove excess oligo before introducing CMG.

#### Unwinding of 10-kb long stretched DNA

15 pM of 10-kb DNA bound to streptavidin on the forked end was introduced in blocking buffer into a flow cell with PEG-biotin-functionalized glass surface and incubated for 1 hour for surface binding. The flow cell was washed with blocking buffer to remove free DNA. Anti-digoxigenin-(anti-dig) coated microspheres (0.05% w/v in blocking buffer) were introduced and incubated for 1 hour for binding to the free digoxigenin-modified end of surface-immobilized DNA molecules. Excess beads were removed by washing the flow cell with blocking buffer. To stretch DNA and attach anti-dig-conjugated beads to the surface, digoxigenin-modified streptavidin (dig-streptavidin, 3 µg/ml) was drawn at 0.2 ml/min in blocking buffer. Free dig-streptavidin was washed out by flushing the flow cell with blocking buffer. CMG-blocking buffer (CMG-binding buffer containing 0.8 mg/ml casein (Sigma Aldrich)) supplemented with 0.3 mM ATPγS was drawn into the flow cell and incubated for 10 minutes. LD655-labelled CMG (15 nM final in CMG-blocking buffer supplemented with 0.3 mM ATPγS) was introduced and incubated for 1 hour. To remove free CMG and initiate DNA unwinding, 20 nM EGFP-RPA was drawn in CMG-blocking buffer containing 3.3 mM ATP, 1% glucose, 0.02 mg/ml glucose oxidase, and 0.04 mg/ml catalase.

### Preparation of anti-digoxigenin-conjugated microspheres

100 μl 5% w/v of 0.45 μm-diameter carboxyl-functionalized polystyrene microspheres (Spherotech, CP-05-10) were washed twice with 0.5 ml coupling buffer (sodium acetate pH 5.0) by centrifugation and resuspended in 0.8 ml coupling buffer. 0.2 mg polyclonal anti-dig antibody (Roche, 11333089001) and 1 mg BSA were dissolved in 0.2 ml coupling buffer and mixed into carboxylated beads. The mixture was incubated 3 hours at room temperature on a rotator. Excess anti-dig and BSA were removed by washing microspheres with 1 ml PBS three consecutive rounds by centrifugation. Microspheres were resuspended in 0.2 ml PBS and stored at 4°C. Before introducing into a flow cell, 2 μl anti-dig microspheres were washed once with 100 μl blocking buffer, resuspended in 100 μl blocking buffer, and briefly sonicated in an ultrasonic water bath (VWR) to break aggregates.

### Preparation of streptavidin-digoxigenin conjugate

10 mg streptavidin (Sigma) was dissolved in sodium bicarbonate pH 8.2. 1 mg digoxigenin-NHS (Sigma, 55865) was dissolved in 100 μl DMSO, mixed into streptavidin solution, and incubated 3 hours at room temperature rotating. Unconjugated digoxenin was removed by exchanging buffer 5 times with 10 ml PBS each round using a spin concentrator (Vivaspin, 20 ml, 30 kDa MWCO). Streptavidin-digoxigenin conjugate was finally concentrated to 3 mg/ml and stored at 4°C.

### Microscope setup and image acquisition

DNA unwinding assays performed with immobilized DNA molecules were imaged on an objective-type TIRF configuration using an inverted microscope (Ti-E, Nikon) equipped with a 100× oil objective (HP Apo TIRF 100xH, N.A.=1.49, Nikon) and automated focus. Fluorescence of Atto647N and LD655 were recorded with excitation wavelength of 647 nm, while Cy3 and EGFP were illuminated with 561-nm and 488-nm lasers, respectively. Images were collected at 100-200 ms exposures per frame on an Andor iXon 897 back-illuminated electron-multiplying CCD camera (Andor Technology).

### Data analysis

Images collected during single-molecule experiments were analysed using NIS-Elements software (Nikon). Images were first aligned to correct the effect of stage drift over time. Bright spots corresponding to Atto647-dye conjugated DNA molecules above a custom threshold observed in the first frame were selected. The fluorescence intensity of each spot at each frame were measured, and exported for further analysis. Reported unwinding efficiencies observed in Fork^ssLag^ and Fork^dsLag^ were measured by dividing the cumulative number molecules that disappeared (i.e. molecules fully unwound) by the total number of molecules present on the first frame. Rate of DNA unwinding on 10-kb linear DNA substrates was measured through the growth rate of EGFP-RPA tracts.

### Fitting and normalization

Fluorescence intensity values on plate reader assays were normalized by fitting the data to Equation 1 (Donmez and Patel, 2008) for integer values of m:

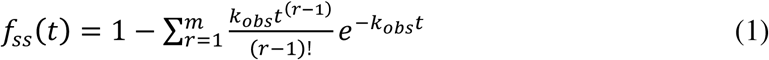

where *fss(t)* is time-dependent extent of DNA unwinding, m is the number of steps, *k_obs_* is the observed unwinding rate and *t* is time. Fluorescence-time traces from CMG unwinding assays were fit using Equation 2 (m=1).

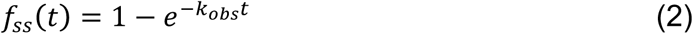

The data from large T antigen- and NS3-mediated DNA unwinding assays were fitted with using Equation 3 (m=2) due to the presence of a lag phase at early time points.

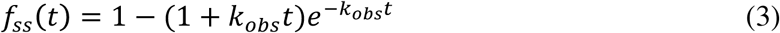

Constant values obtained subsequent to fitting were used to normalize unwinding signals. Normalized data were averaged and plotted against time.

### Statistical analysis

Throughout the manuscript, the data are represented as average ± standard deviation of multiple experiments. Prism (GraphPad Software, La Jolla, CA, USA) was used to plot all graphs presented and for statistical analysis in this study. ImageJ was used to quantify band intensities in gel images.

## Acknowledgements

We thank Daniel Burnham and Alessandro Costa for critical reading of the manuscript, and Peptide Chemistry and Cell Services science technology platforms at the Francis Crick Institute for providing peptides. Additional thanks to Mauro Modesti for RPA and EGFP-RPA expression vectors, Charles Rice for NS3 expression vector, and David Liu for Sortase expression vector. This work was supported by the Francis Crick Institute, which receives its core funding from Cancer Research UK (FC001221), the UK Medical Research Council (FC001221), and the Wellcome Trust (FC001221). G.C. was supported by a PhD fellowship from the Boehringer Ingelheim Fonds.

## Author Contributions

H.B.K and H.Y. designed the study and conducted experiments. S.X. and G.C. expressed and purified LD655-labelled CMG. M.S.S. expressed and purified Sortase. H.B.K. purified all other recombinant proteins. H.B.K and H.Y. interpreted the data and wrote the paper with input from other authors.

---

**Supplementary Figure 1.**
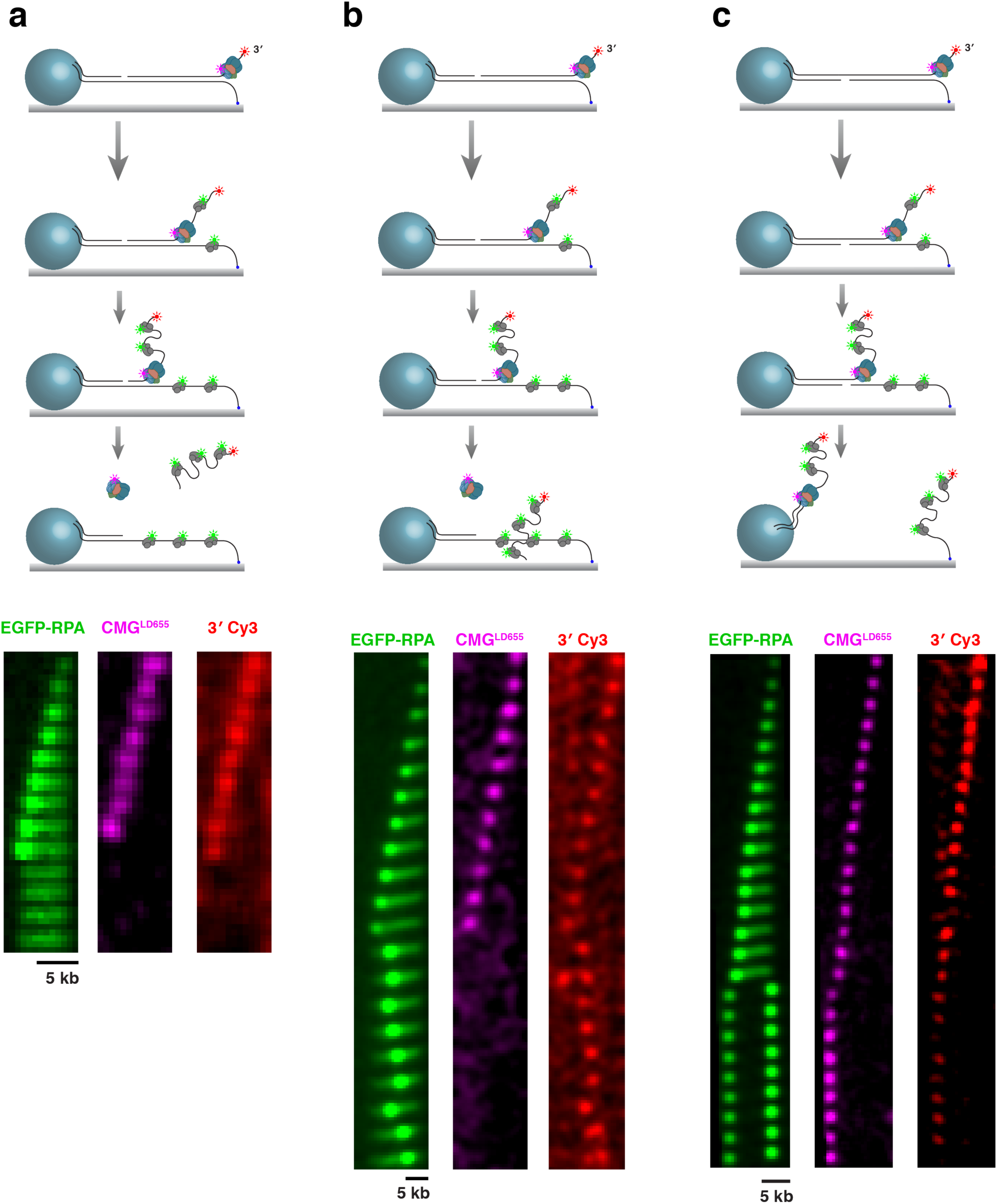
Single-molecule visualization of processive DNA unwinding by CMG. **a-b**, Cartoon depictions of CMG meeting a nick on the leading-strand template. The leading-strand template either dissociates from (**a**) or diffuses along (**b**) the surface-immobilized DNA. Bottom images show corresponding example events. CMG does not remain on the translocation strand in **b** suggesting that it runs off the free 5’ end. **c,** CMG collides with a lagging-strand nick resulting in breakage of DNA. CMG remains on DNA. Bottom images show an example event for CMG encountering a lagging-strand nick.

**Supplementary Figure 2.**
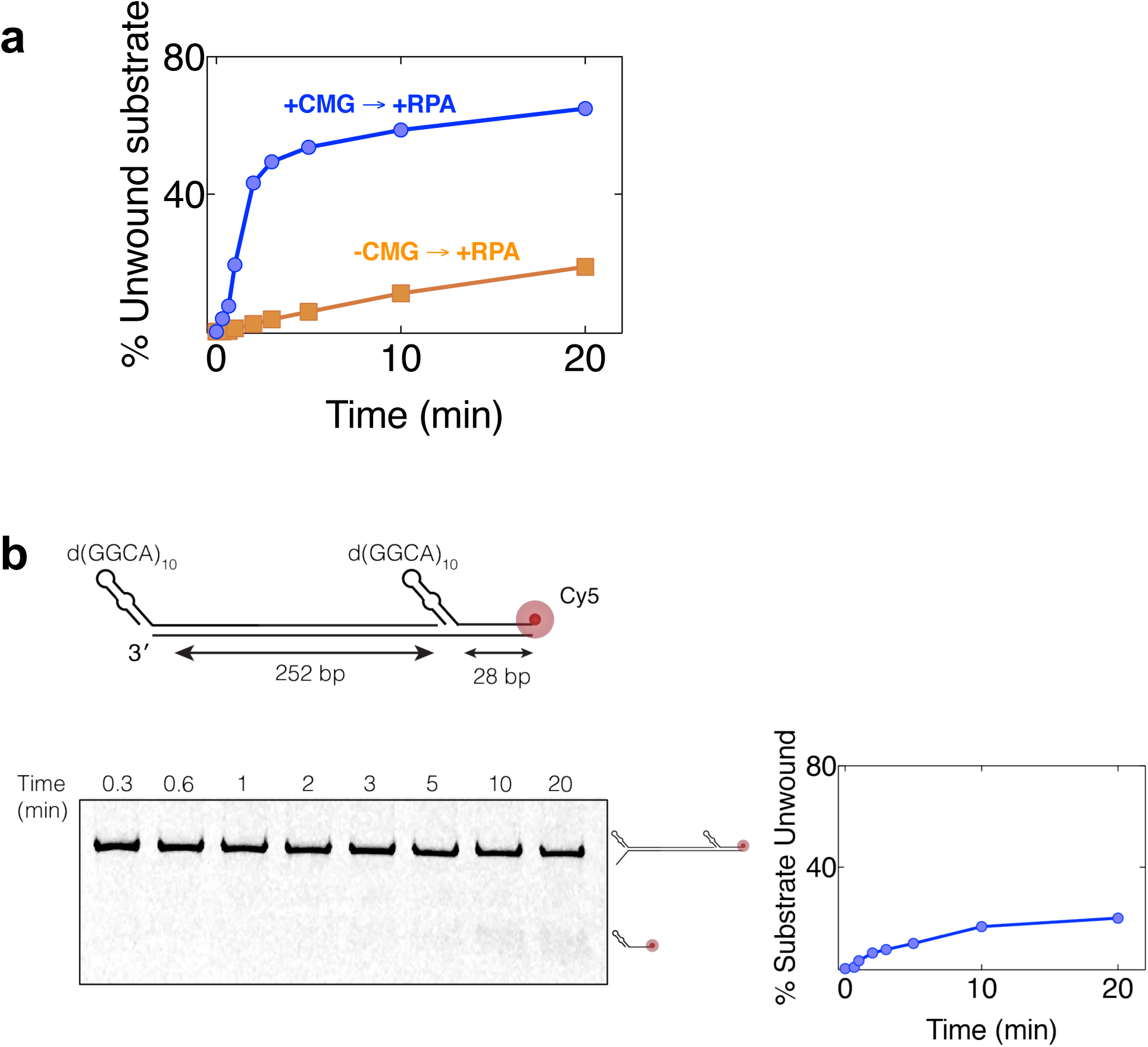
Displacement of the Cy5-modified oligonucleotide downstream of 252-bp dsDNA is dependent on CMG binding to the upstream 3’ dT_40_ ssDNA. **a,** The same fork DNA used in Figure 2c was incubated in buffer containing or lacking CMG. RPA was then added together with ATP. RPA displaced Cy5-labelled strand to some extent even in the absence of CMG. The percentage of DNA unwound by RPA alone (-CMG→+RPA) was subtracted from that by CMG and RPA (+CMG→+RPA) to determine CMG-mediated unwinding in the presence of RPA. The data plotted in Figure 2e demonstrates measurements after this correction. **b,** The DNA substrate containing 252-bp long dsDNA, followed by 28-bp duplex and Cy5-modification on the excluded strand was incubated with CMG in the presence of ATPγS. This substrate lacked 3’ dT_40_ ssDNA overhang. Following CMG incubation with DNA, ATP and RPA were added, and the reaction was further incubated at 30°C for indicated periods of time. The right panel demonstrates the percentage of strand displacement displaced as a function of time as quantified from the gel. Cy5-modified strand was displaced to the same degre as the reaction lacking CMG in Figure S2A (-CMG→+RPA) indicating that CMG binding to 3’ dT_40_ upstream of the 252-bp dsDNA was needed for CMG-mediated unwinding.

**Supplementary Figure 3.**
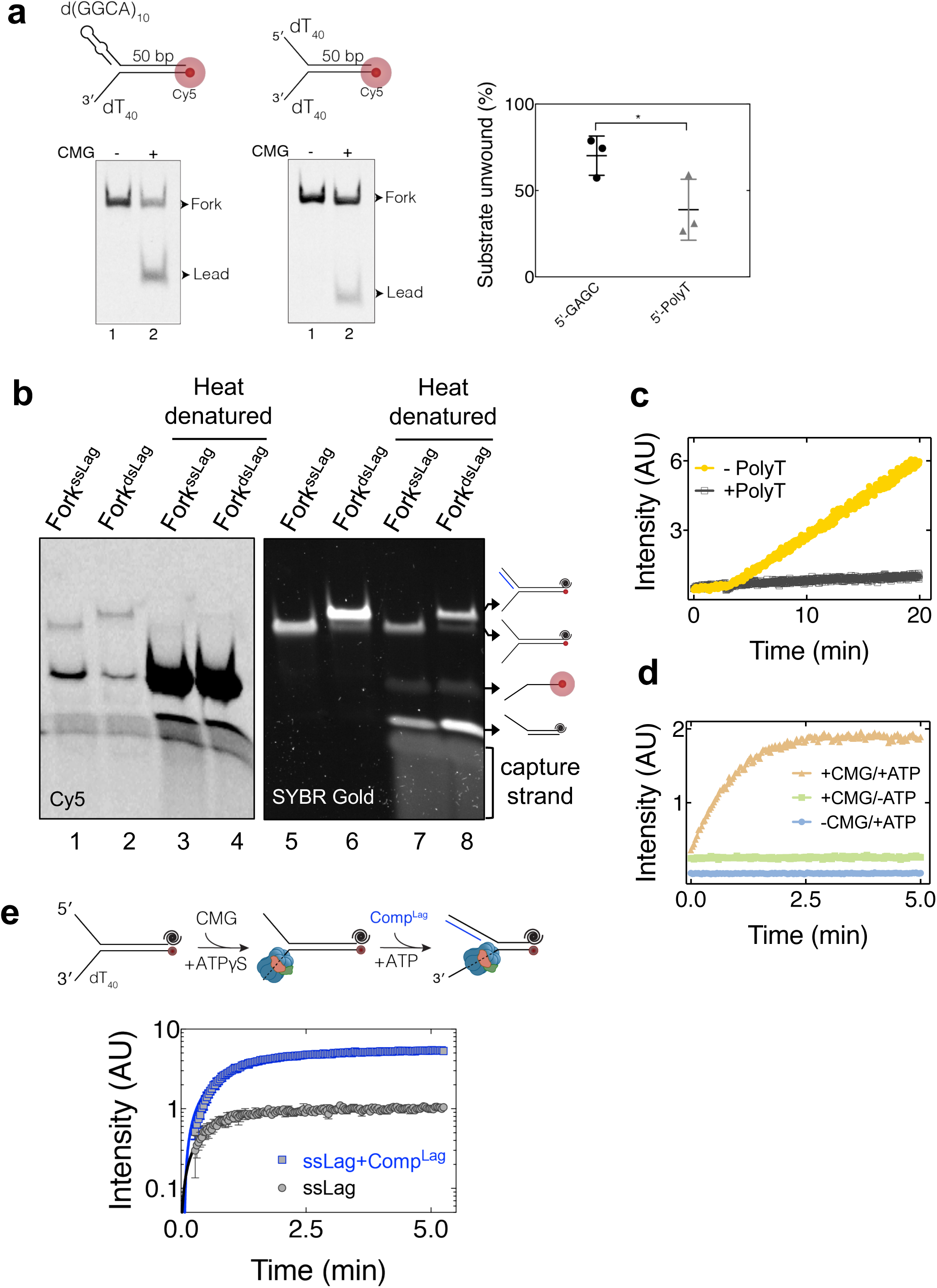
The impact of the lagging-strand arm on CMG helicase activity. **a,**Fork DNA containing 3’ dT_40_, 50-bp duplex, 5’ leading-strand Cy5, and either dT_40_ or d(GAGC)_10_ lagging-strand arm was unwound by CMG and separated on 8% PAGE. The right panel shows quantification of DNA unwound (mean ± SD, n=3). **b,** Cy5/BHQ2-labelled fork DNA containing either single-stranded (Fork^ssLag^) or double-stranded (Fork^dsLag^) lagging-strand arm was separated on 8% PAGE. Fluorescence of Cy5-labelled strand is quenched and increases upon heat denaturation (compare lanes, 1-2 and 3-4). The right panel shows the same gel visualized under UV after SYBR Gold staining. A capture strand complementary to the BHQ2-modified strand was included in heat-denatured samples to prevent reannealing. **c,** CMG was mixed with ATP and Cy5/BHQ2-labelled Fork^dsLag^ in the absence (grey) or presence (yellow) of free dT_40_ oligonucleotide. Excess dT_40_ prevents CMG binding to fork DNA and subsequent unwinding. **d,** Time-dependent fluorescence intensity of Cy5/BHQ2-labelled Fork^dsLag^ containing 28-bp parental duplex. Fluorescence increase is strictly dependent on the presence of CMG during ATPγS incubation as well as subsequent ATP addition. **e,** Cy5/BHQ2-labelled Fork^ssLag^ was pre-incubated with CMG. ATP solution with (blue) or without (black) an oligonucleotide complementary to the 22-nt lagging-strand arm (Comp^Lag^) was added, and fluorescence intensity was measured.

**Supplementary Figure 4.**
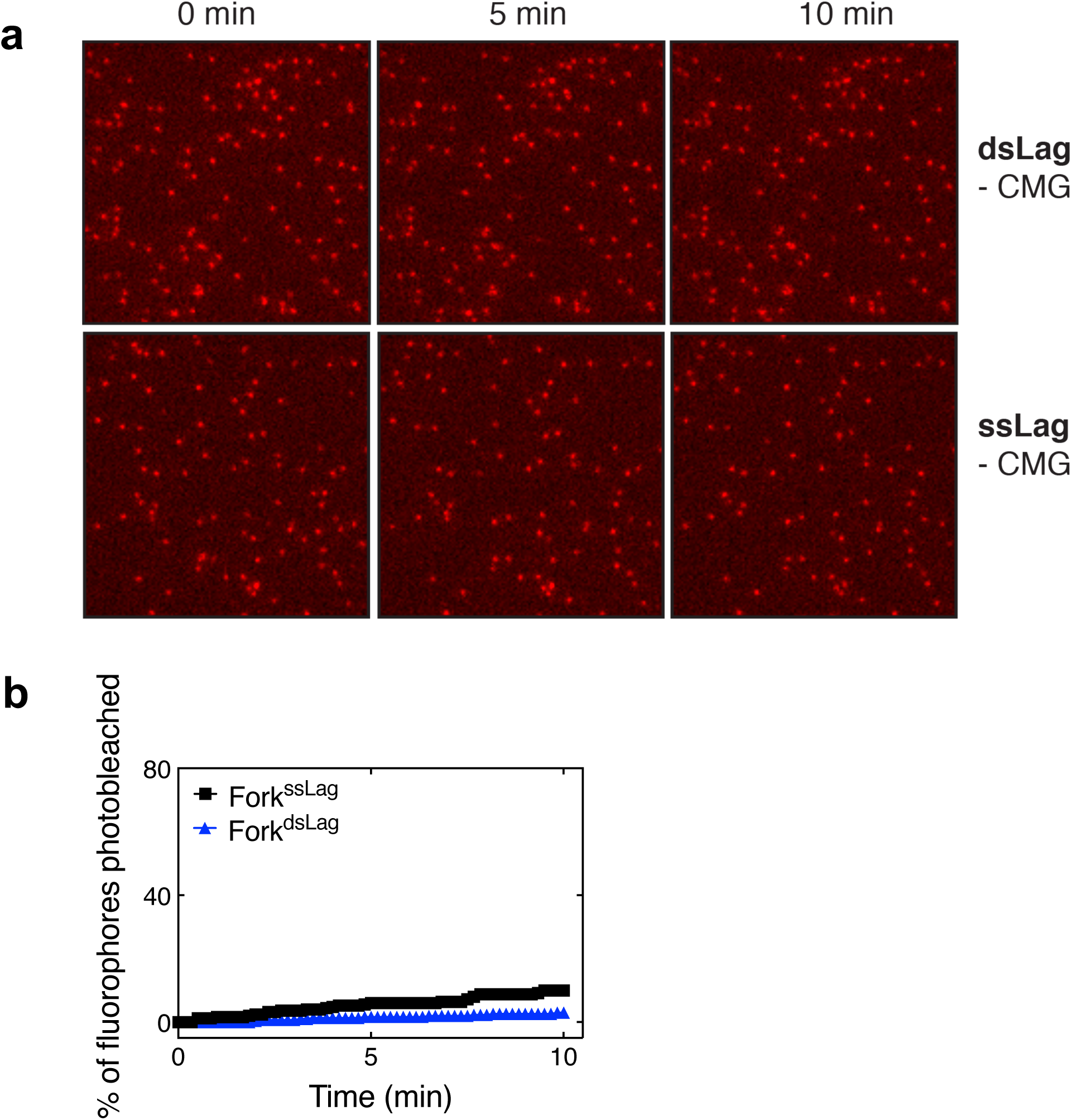
Photostability of Atto647N fluorophore. **a,** Atto647N-labelled fork DNA was immobilized and imaged under the same experimental conditions for DNA unwinding assay (Figure 4) except CMG was omitted from ATPγS buffer. **b,** Percentage of Atto647N spots photobleached as a function of time (Fork^ssLag^ N=251 molecules, Fork^dsLag^ N=306 molecules, each from two independent experiments).

**Supplementary Figure 5.**
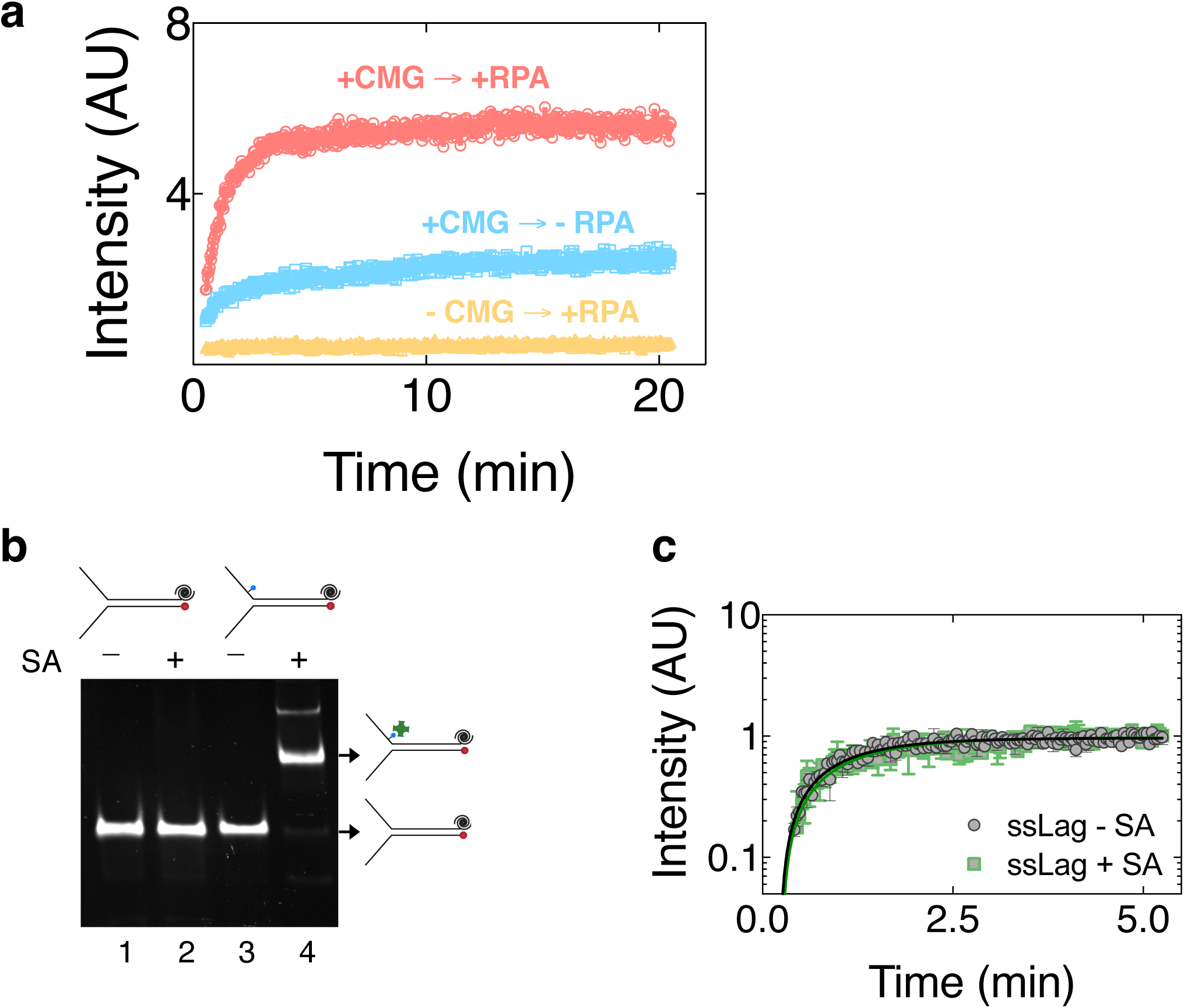
Binding of a protein to the lagging-strand arm of fork DNA. **a,** Fork^ssLag^ was incubated with buffer containing or lacking CMG in the presence of ATPγS for 90 minutes. Buffer or RPA was subsequently added and incubated for 10 minutes. ATP was then added to initiate CMG translocation. A competitor oligonucleotide was included with ATP addition to prevent further RPA and CMG binding during unwinding. When CMG was pre-bound to the fork, addition of RPA stimulated unwinding (compare +CMG→+RPA to +CMG→-RPA) similar to shown in Figure 5. Importantly, when CMG was omitted from the reaction, addition of RPA did not lead to detectable unwinding (-CMG→+RPA). **b,** Binding of streptavidin (SA) to the lagging-strand biotin on fork DNA is monitored by separating DNA on 8% PAGE and subsequent staining with SYBR Gold. Addition of streptavidin to biotin-modified fork leads to mobility shift (lane 4). **c,** CMG-catalyzed single turn-over unwinding of fork DNA lacking a biotin on the lagging-strand arm in the absence (black) and presence (green) of streptavidin (SA). Data represent mean ± SD (n=3).

**Supplementary Figure 6.**
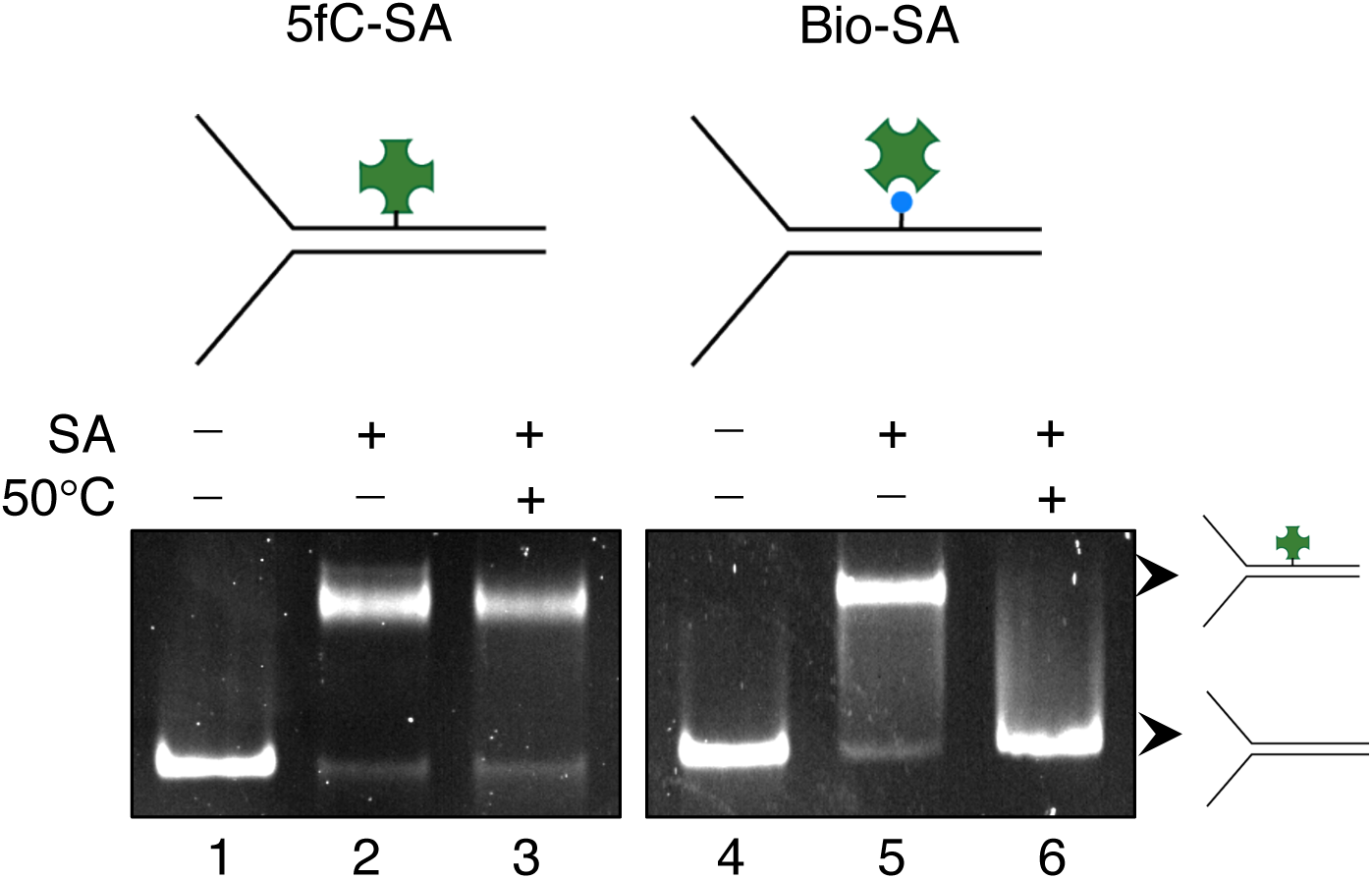
5-Formyl-cystosine-streptavidin crosslinked fork DNA substrate. Fork DNA containing either 5-formyl-cytosine-streptavidin (5fC-SA) crosslink (lane 2) or biotin-streptavidin (bio-SA) complex (lane 4) was heated to 50°C for 10 minutes in the presence of 1 μM 5fC-modified oligonucleotide or 1 μM free biotin, respectively, and separated on 8% PAGE. While streptavidin dissociated from biotin upon heat treatment (lane 6), 5fC-SA crosslink remained stable (lane 3).

**Supplementary Table 1.** Oligonucleotides used in each DNA substrate.

| **Substrate Name** | **Oligonucleotides Used** | **Associated Figures** |
| --- | --- | --- |
| 236-bp fork template<br>(internally Cy5 labelled) | Oligo-2, Oligo-3, Oligo-4, Oligo-5 | Fig. 2a |
| 252-bp fork template<br>(3'-Cy5) | Oligo-2, Oligo-17, Oligo-4, Oligo-5, Oligo-15,<br>Oligo-16, Oligo-Cy5-6 | Fig. 2c-d,<br>Supp. Fig. 2a |
| 252-bp fork template<br>lacking 3'-PolyT tail | Oligo-2, Oligo-3, Oligo-4, Oligo-18, Oligo-15,<br>Oligo-16, Oligo-Cy5-6 | Supp. Fig. 2b |
| Fork<sup>ssLag</sup> | Oligo-BHQ2-2, Oligo-Cy5-4 | Fig. 3, 5a, 6a,<br>Supp. Fig. 3e |
| Fork<sup>dsLag</sup> | Oligo-12, Oligo-BHQ2-2, Oligo-Cy5-4 | Fig. 3, 5b,<br>Supp. Fig. 3b-d |
| Fork<sup>ssLag</sup>-Bio | Oligo-BHQ2-Bio, Oligo-Cy5-4 | Fig. 5c, Supp.<br>Fig. 5b |
| Fork-5'-PolyT | Oligo-Bio-2, Oligo-Cy5-5 | Supp. Fig. 3a |
| Fork-5'-GAGC | Oligo-Bio-1, Oligo-Cy5-5 | Supp. Fig. 3a |
| Fork<sup>ssLag</sup>-TIRF | Oligo-Atto, Oligo-19, Oligo-Bio-5 | Fig. 4, Supp.<br>Fig. 4 |
| Fork<sup>dsLag</sup>-TIRF | Oligo-Atto, Oligo-19, Oligo-Bio-5, Oligo-12 | Fig. 4, Supp.<br>Fig. 4 |
| 60-bp fork with 5'-<br>GAGC | Oligo-11, Oligo-BHQ2-1, Oligo-Cy5-1 | Fig. 6b |
| Fork (5fC-SA<br>crosslink) | Oligo-5fC-1, Oligo-9, Oligo-14, Oligo-IBRQ | Fig. 7 |
| Fork (no crosslink) | Oligo-7, Oligo-14, Oligo-Cy5-2, Oligo-IBRQ | Fig. 7 |
| Fork-Bio-SA | Oligo-10, Oligo-Bio-3, Oligo-Cy5-3 | Supp. Fig. 6 |
| 10kb-substrate (TIRF) | pUC19 as the PCR template, Oligo-Bio-4,<br>Oligo-Cy3, Oligo-1, Oligo-8 | Fig. 1, Supp.<br>Fig. 1 |

**Supplementary Table 2.**
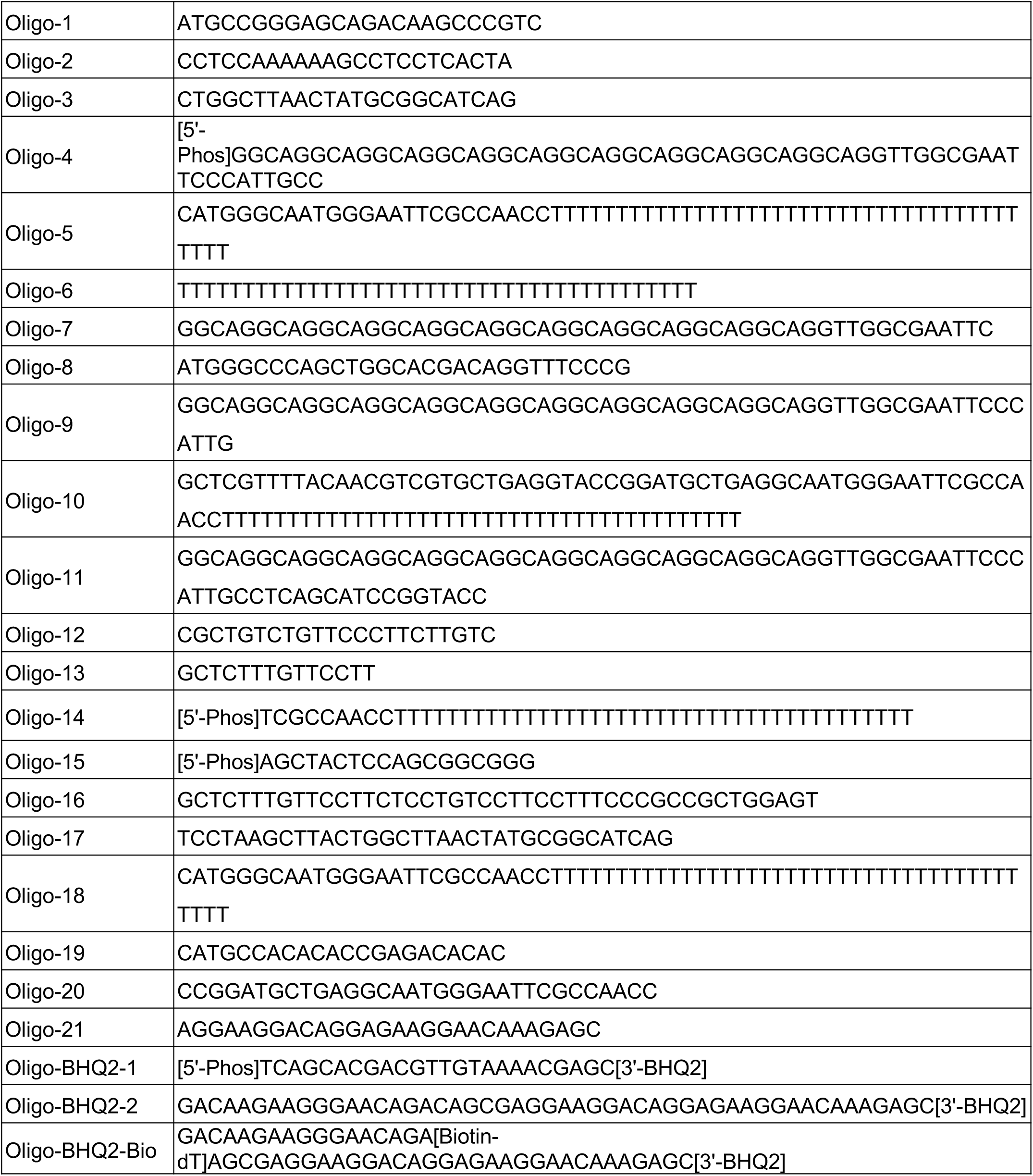
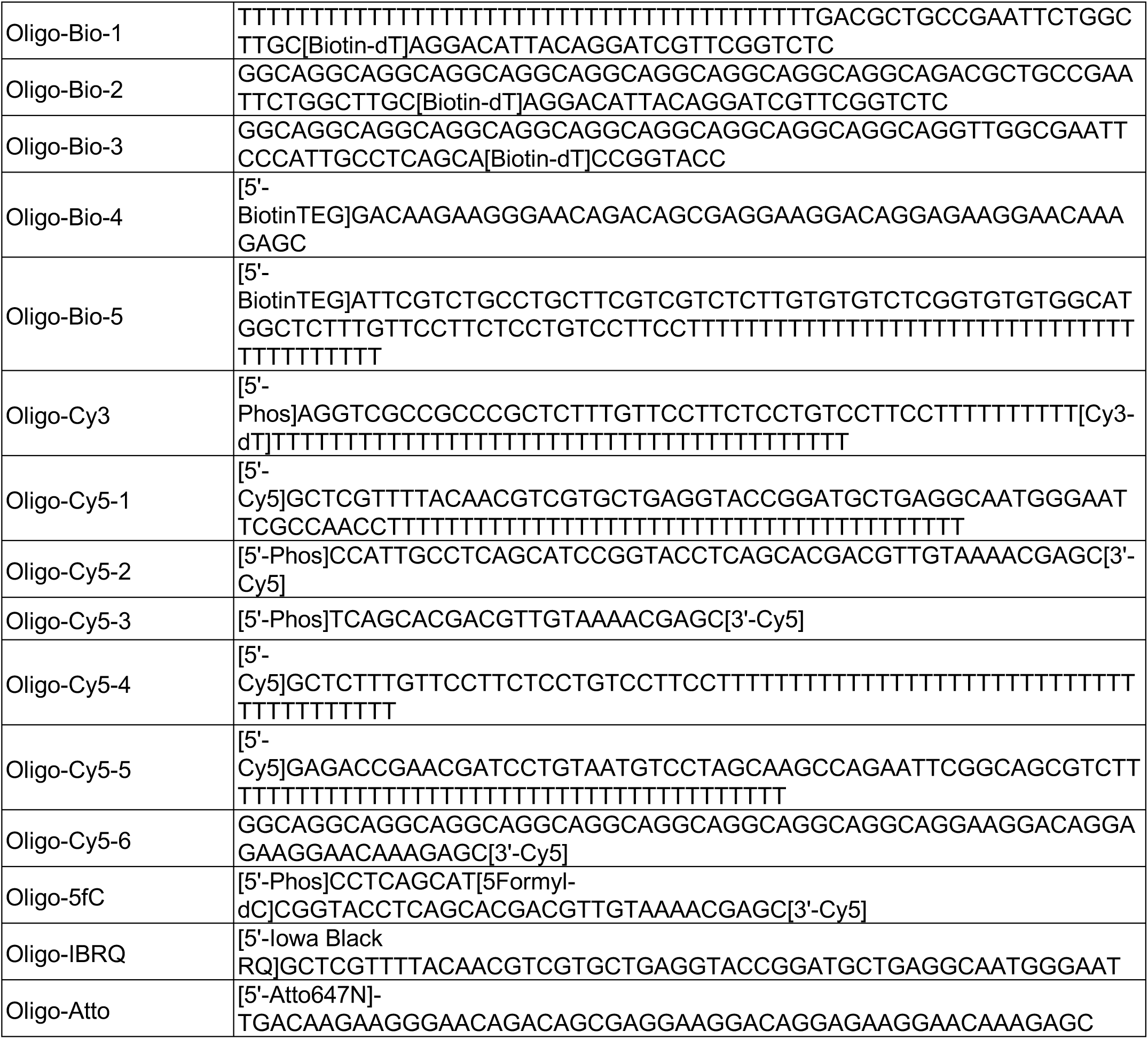
Sequences of oligonucleotides used to prepare DNA substrates.

## References

Abid Ali, F., Douglas, M.E., Locke, J., Pye, V.E., Nans, A., Diffley, J.F.X., and Costa, A. (2017). Cryo-EM structure of a licensed DNA replication origin. Nat. Commun. 8, 2241.

Anglana, M., Apiou, F., Bensimon, A., and Debatisse, M. (2003). Dynamics of DNA Replication in Mammalian Somatic Cells: Nucleotide Pool Modulates Origin Choice and Interorigin Spacing. Cell 114, 385–394.

Arias, E.E., and Walter, J.C. (2007). Strength in numbers: preventing rereplication via multiple mechanisms in eukaryotic cells. Genes Dev. 21, 497–518.

Atkinson, J., Gupta, M.K., and McGlynn, P. (2010). Interaction of Rep and DnaB on DNA. Nucleic Acids Res. 39, 1351–1359.

Blumenthal, A.B., Kriegstein, H.J., and Hogness, D.S. (1974). The Units of DNA Replication in Drosophila melanogaster Chromosomes. Cold Spring Harbor Symp. Quant. Biol. 38, 205–223.

Burnham, D.R., Kose, H.B., Hoyle, R.B., and Yardimci, H. (2019). The mechanism of DNA unwinding by the eukaryotic replicative helicase. Nat. Commun. 10, 2159.

Carney, S.M., Gomathinayagam, S., Leuba, S.H., and Trakselis, M.A. (2017). Bacterial DnaB helicase interacts with the excluded strand to regulate unwinding. J. Biol. Chem. 292, 19001–19012.

Chen, I., Dorr, B.M., and Liu, D.R. (2011). A general strategy for the evolution of bond-forming enzymes using yeast display. Proc. Natl. Acad. Sci. USA 108, 11399–11404.

Donmez, I., and Patel, S.S. (2008). Coupling of DNA unwinding to nucleotide hydrolysis in a ring-shaped helicase. EMBO J. 27, 1718–1726.

Douglas, M.E., Ali, F.A., Costa, A., and Diffley, J.F.X. (2018). The mechanism of eukaryotic CMG helicase activation. Nature 555, 265–268.

Duxin, J.P., Dewar, J.M., Yardimci, H., and Walter, J.C. (2014). Repair of a DNA-protein crosslink by replication-coupled proteolysis. Cell 159, 346–357.

Egelman, E.H., Yu, X., Wild, R., Hingorani, M.M., and Patel, S.S. (1995). Bacteriophage T7 helicase/primase proteins form rings around single-stranded DNA that suggest a general structure for hexameric helicases. Proc. Natl. Acad. Sci. USA 92, 3869–3873.

Eickhoff, P., Kose, H.B., Martino, F., Petojevic, T., Abid Ali, F., Locke, J., Tamberg, N., Nans, A., Berger, J.M., Botchan, M.R., et al. (2019). Molecular Basis for ATP-Hydrolysis-Driven DNA Translocation by the CMG Helicase of the Eukaryotic Replisome. Cell Rep. 28, 2673–2688.

Evrin, C., Clarke, P., Zech, J., Lurz, R., Sun, J., Uhle, S., Li, H., Stillman, B., and Speck, C. (2009). A double-hexameric MCM2-7 complex is loaded onto origin DNA during licensing of eukaryotic DNA replication. Proc. Natl. Acad. Sci. USA 106, 20240–20245.

Fu, Y.V., Yardimci, H., Long, D.T., Ho, T.V., Guainazzi, A., Bermudez, V.P., Hurwitz, J., van Oijen, A., Schärer, O.D., and Walter, J.C. (2011). Selective bypass of a lagging strand roadblock by the eukaryotic replicative DNA helicase. Cell 146, 931–941.

Gai, D., Wang, D., Li, S.-X., and Chen, X.S. (2016). The structure of SV40 large T hexameric helicase in complex with AT-rich origin DNA. eLife 5, e18129.

Gao, Y., Cui, Y., Fox, T., Lin, S., Wang, H., de Val, N., Zhou, Z.H., and Yang, W. (2019). Structures and operating principles of the replisome. Science 363, eaav7003.

Georgescu, R., Yuan, Z., Bai, L., de Luna Almeida Santos, R., Sun, J., Zhang, D., Yurieva, O., Li, H., and O’Donnell, M.E. (2017). Structure of eukaryotic CMG helicase at a replication fork and implications to replisome architecture and origin initiation. Proc. Natl. Acad. Sci. USA 114, E697–E706.

Graham, B.W., Schauer, G.D., Leuba, S.H., and Trakselis, M.A. (2011). Steric exclusion and wrapping of the excluded DNA strand occurs along discrete external binding paths during MCM helicase unwinding. Nucleic Acids Res. 39, 6585–6595.

Gu, M., and Rice, C.M. (2010). Three conformational snapshots of the hepatitis C virus NS3 helicase reveal a ratchet translocation mechanism. Proc. Natl. Acad. Sci. USA 107, 521–528.

Hizume, K., Endo, S., Muramatsu, S., Kobayashi, T., and Araki, H. (2018). DNA polymerase ε-dependent modulation of the pausing property of the CMG helicase at the barrier. Genes Dev. 32, 1315–1320.

Ilves, I., Petojevic, T., Pesavento, J.J., and Botchan, M.R. (2010). Activation of the MCM2-7 Helicase by Association with Cdc45 and GINS Proteins. Mol. Cell 37, 247–258.

Jeong, Y.-J., Rajagopal, V., and Patel, S.S. (2013). Switching from single-stranded to double-stranded DNA limits the unwinding processivity of ring-shaped T7 DNA helicase. Nucleic Acids Res. 41, 4219–4229.

Ji, S., Shao, H., Han, Q., Seiler, C.L., and Tretyakova, N.Y. (2017). Reversible DNA–Protein Cross-Linking at Epigenetic DNA Marks. Angew. Chem., Int. Ed. 56, 14130–14134.

Johnson, D.S., Bai, L., Smith, B.Y., Patel, S.S., and Wang, M.D. (2007). Single-Molecule Studies Reveal Dynamics of DNA Unwinding by the Ring-Shaped T7 Helicase. Cell 129, 1299–1309.

Kang, Y.-H., Galal, W.C., Farina, A., Tappin, I., and Hurwitz, J. (2012). Properties of the human Cdc45/Mcm2-7/GINS helicase complex and its action with DNA polymerase ε in rolling circle DNA synthesis. Proc. Natl. Acad. Sci. USA 109, 6042–6047.

Kaplan, D.L. (2000). The 3′-tail of a forked-duplex sterically determines whether one or two DNA strands pass through the central channel of a replication-fork helicase. J. Mol. Biol. 301, 285–299.

Kaplan, D.L., Davey, M.J., and O’Donnell, M. (2003). Mcm4,6,7 Uses a “Pump in Ring” Mechanism to Unwind DNA by Steric Exclusion and Actively Translocate along a Duplex. J. Biol. Chem. 278, 49171–49182.

Kemmerich, F.E., Daldrop, P., Pinto, C., Levikova, M., Cejka, P., and Seidel, R. (2016). Force regulated dynamics of RPA on a DNA fork. Nucleic Acids Res. 44, 5837–5848.

Kim, S., Dallmann, H.G., McHenry, C.S., and Marians, K.J. (1996). Coupling of a Replicative Polymerase and Helicase: A τ–DnaB Interaction Mediates Rapid Replication Fork Movement. Cell 84, 643–650.

Klaue, D. (2012). DNA Unwinding by Helicases Investigated on the Single Molecule Level. In Institut fur Biophysik (Technische Universität Dresden).

Kose, H.B., Larsen, N.B., Duxin, J.P., and Yardimci, H. (2019). Dynamics of the Eukaryotic Replicative Helicase at Lagging-Strand Protein Barriers Support the Steric Exclusion Model. Cell Rep. 26, 2113–2125.

Langston, L., and O’Donnell, M. (2017). Action of CMG with strand-specific DNA blocks supports an internal unwinding mode for the eukaryotic replicative helicase. eLife 6, e23449.

Langston, L.D., Zhang, D., Yurieva, O., Georgescu, R.E., Finkelstein, J., Yao, N.Y., Indiani, C., and O’Donnell, M.E. (2014). CMG helicase and DNA polymerase ε form a functional 15-subunit holoenzyme for eukaryotic leading-strand DNA replication. Proc. Natl. Acad. Sci. USA 111, 15390–15395.

Lee, S.-J., Syed, S., Enemark, E.J., Schuck, S., Stenlund, A., Ha, T., and Joshua-Tor, L. (2014). Dynamic look at DNA unwinding by a replicative helicase. Proc. Natl. Acad. Sci. USA 111, E827–E835.

Lewis, J.S., Spenkelink, L.M., Schauer, G.D., Hill, F.R., Georgescu, R.E., O’Donnell, M.E., and van Oijen, A.M. (2017). Single-molecule visualization of Saccharomyces cerevisiae leading-strand synthesis reveals dynamic interaction between MTC and the replisome. Proc. Natl. Acad. Sci. USA 114, 10630–10635.

Li, F., Zhang, Y., Bai, J., Greenberg, M.M., Xi, Z., and Zhou, C. (2017). 5-Formylcytosine Yields DNA–Protein Cross-Links in Nucleosome Core Particles. J. Am. Chem. Soc. 139, 10617–10620.

Li, H., O’Donnell, M.E. (2018). The eukaryotic CMG helicase at the replication fork: Emerging architecture reveals an unexpected mechanism. Bioessays 40.

Lionnet, T., Spiering, M.M., Benkovic, S.J., Bensimon, D., and Croquette, V. (2007). Real-time observation of bacteriophage T4 gp41 helicase reveals an unwinding mechanism. Proc. Natl. Acad. Sci. USA 104, 19790–19795.

Moyer, S.E., Lewis, P.W., and Botchan, M.R. (2006). Isolation of the Cdc45/Mcm2–7/GINS (CMG) complex, a candidate for the eukaryotic DNA replication fork helicase. Proc. Natl. Acad. Sci. USA 103, 10236–10241.

Neelsen, K.J., and Lopes, M. (2015). Replication fork reversal in eukaryotes: from dead end to dynamic response. Nat. Rev. Mol. Cell Biol. 16, 207–220.

Noguchi, Y., Yuan, Z., Bai, L., Schneider, S., Zhao, G., Stillman, B., Speck, C., and Li, H. (2017). Cryo-EM structure of Mcm2-7 double hexamer on DNA suggests a lagging-strand DNA extrusion model. Proc. Natl. Acad. Sci. USA 114, E9529–E9538.

Pang, P.S., Jankowsky, E., Planet, P.J., and Pyle, A.M. (2002). The hepatitis C viral NS3 protein is a processive DNA helicase with cofactor enhanced RNA unwinding. EMBO J. 21, 1168–1176.

Petojevic, T., Pesavento, J.J., Costa, A., Liang, J., Wang, Z., Berger, J.M., and Botchan, M.R. (2015). Cdc45 (cell division cycle protein 45) guards the gate of the Eukaryote Replisome helicase stabilizing leading strand engagement. Proc. Natl. Acad. Sci. USA 112, E249–E258.

Raghuraman, M.K., Winzeler, E.A., Collingwood, D., Hunt, S., Wodicka, L., Conway, A., Lockhart, D.J., Davis, R.W., Brewer, B.J., and Fangman, W.L. (2001). Replication Dynamics of the Yeast Genome. Science 294, 115–121.

Remus, D., Beuron, F., Tolun, G., Griffith, J.D., Morris, E.P., and Diffley, J.F.X. (2009). Concerted loading of Mcm2–7 double hexamers around DNA during DNA replication origin licensing. Cell 139, 719–730.

Ribeck, N., Kaplan, D.L., Bruck, I., and Saleh, O.A. (2010). DnaB Helicase Activity Is Modulated by DNA Geometry and Force. Biophys. J. 99, 2170–2179.

Shin, J.-H., Jiang, Y., Grabowski, B., Hurwitz, J., and Kelman, Z. (2003). Substrate Requirements for Duplex DNA Translocation by the Eukaryal and Archaeal Minichromosome Maintenance Helicases. J. Biol. Chem. 278, 49053–49062.

Sparks, J.L., Chistol, G., Gao, A.O., Räschle, M., Larsen, N.B., Mann, M., Duxin, J.P., and Walter, J.C. (2019). The CMG Helicase Bypasses DNA-Protein Cross-Links to Facilitate Their Repair. Cell 176, 167–181.

Stano, N.M., Jeong, Y.J., Donmez, I., Tummalapalli, P., Levin, M.K., and Patel, S.S. (2005). DNA synthesis provides the driving force to accelerate DNA unwinding by a helicase. Nature 435, 370–373.

Strycharska, M.S., Arias-Palomo, E., Lyubimov, A.Y., Erzberger, J.P., O’Shea,V.L., Bustamante, C.J., and Berger, J.M. (2013). Nucleotide and partner-pro-tein control of bacterial replicative helicase structure and function. Mol. Cell. 52, 844–854.

Syed, S., Pandey, M., Patel, Smita S., and Ha, T. (2014). Single-Molecule Fluorescence Reveals the Unwinding Stepping Mechanism of Replicative Helicase. Cell Rep. 6, 1037–1045.

Taylor, M.R.G., and Yeeles, J.T.P. (2019). Dynamics of Replication Fork Progression Following Helicase–Polymerase Uncoupling in Eukaryotes. J. Mol. Biol. 431, 2040–2049.

Trakselis, M. (2016). Structural Mechanisms of Hexameric Helicase Loading, Assembly, and Unwinding. F1000Research 5, 111.

Wasserman, M.R., Schauer, G.D., O’Donnell, M.E., and Liu, S. (2019). Replication Fork Activation Is Enabled by a Single-Stranded DNA Gate in CMG Helicase. Cell 178, 600–611.

Wiekowski, M., Schwarz, M.W., and Stahl, H. (1988). Simian virus 40 large T antigen DNA helicase. Characterization of the ATPase-dependent DNA unwinding activity and its substrate requirements. J. Biol. Chem. 263, 436–442.

Yardimci, H., Loveland, A.B., van Oijen, A.M., and Walter, J.C. (2012a). Single-molecule analysis of DNA replication in Xenopus egg extracts. Methods 57, 179–186.

Yardimci, H., Wang, X., Loveland, A.B., Tappin, I., Rudner, D.Z., Hurwitz, J., van Oijen, A.M., and Walter, J.C. (2012b). Bypass of a protein barrier by a replicative DNA helicase. Nature 492, 205–209.

Yeeles, J.T.P., Janska, A., Early, A., and Diffley, J.F.X. (2017). How the Eukaryotic Replisome Achieves Rapid and Efficient DNA Replication. Mol. Cell 65, 105–116.

